# Structures of the ATP-fueled ClpXP proteolytic machine bound to protein substrate

**DOI:** 10.1101/704999

**Authors:** Xue Fei, Tristan A. Bell, Simon Jenni, Benjamin M. Stinson, Tania A. Baker, Stephen C. Harrison, Robert T. Sauer

## Abstract

ClpXP is an ATP-dependent protease in which the ClpX AAA+ motor binds, unfolds, and translocates specific protein substrates into the degradation chamber of ClpP. We present cryo-EM studies of the *E. coli* enzyme that show how asymmetric hexameric rings of ClpX bind symmetric heptameric rings of ClpP and interact with protein substrates. Subunits in the ClpX hexamer assume a spiral conformation and interact with two-residue segments of substrate in the axial channel, as observed for other AAA+ proteases and protein-remodeling machines. Strictly sequential models of ATP hydrolysis and a power stroke that moves two residues of the substrate per translocation step have been inferred from these structural features for other AAA+ unfoldases, but biochemical and single-molecule biophysical studies indicate that ClpXP operates by a probabilistic mechanism in which five to eight residues are translocated for each ATP hydrolyzed. We propose structure-based models that could account for the functional results.

## INTRODUCTION

AAA+ motors harness the energy of ATP hydrolysis to carry out mechanical tasks in cells (Erzberger and Berger, 2006). In the ClpXP protease, for example, AAA+ ClpX ring hexamers bind target proteins, unfold them, and translocate the unfolded polypeptide through an axial channel and into the peptidase chamber of ClpP, which consists of two heptameric rings (Figure 1A; Wang et al., 1997; Grimaud et al., 1998; Ortega et al., 2002; Sauer and Baker, 2011). In the absence of ClpX or another AAA+ partner, small peptides diffuse into the ClpP chamber through narrow axial pores, but larger peptides and native proteins are excluded and escape degradation (Grimaud et al., 1998; Lee et al., 2010a). The substrates of *Escherichia coli* ClpXP include aberrant ssrA-tagged proteins, produced by abortive translation, and normal cellular proteins synthesized with degradation tags that program rapid turnover (Baker and Sauer, 2012; Keiler, 2015).

**Figure 1.**
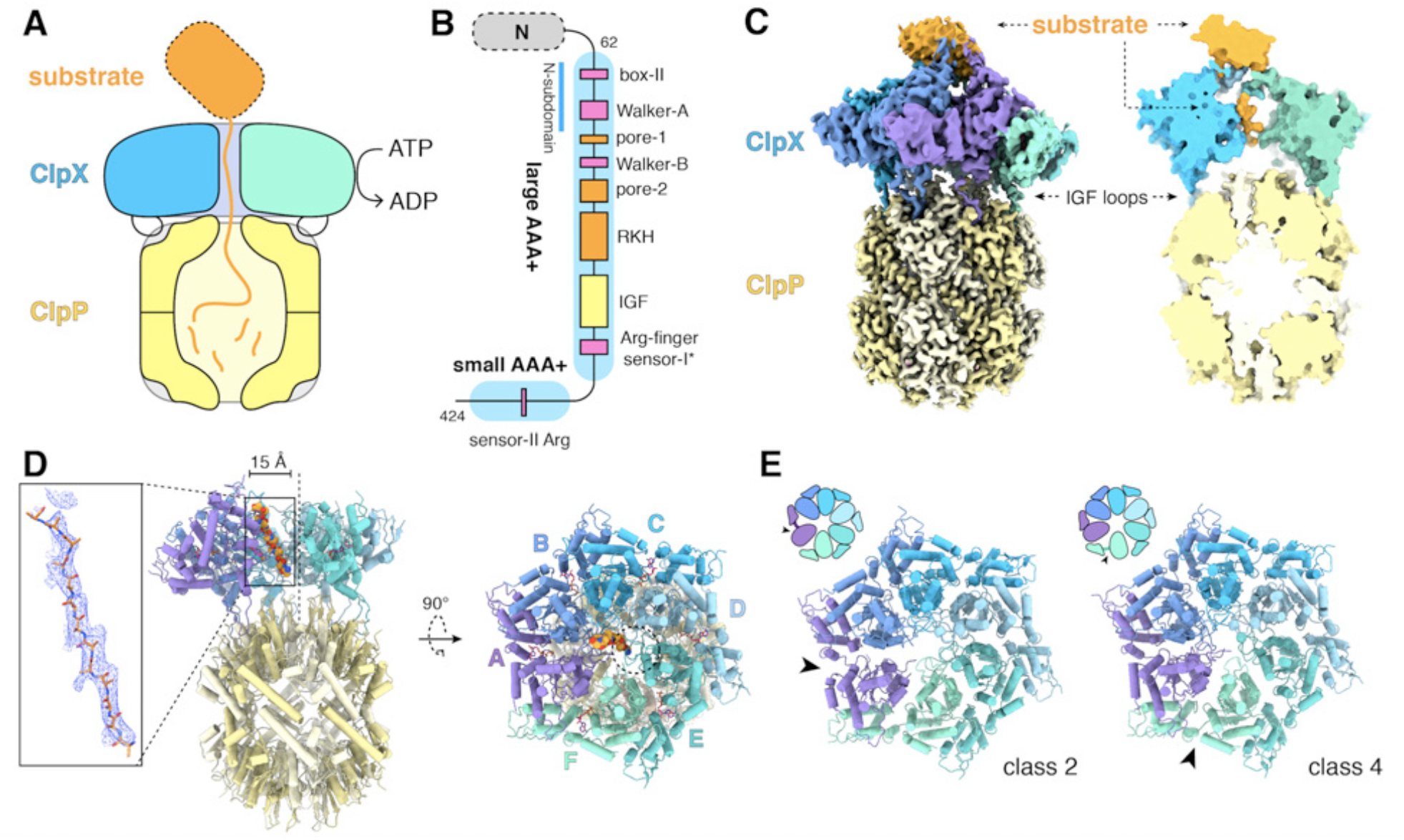
ClpXP protease. (A) Schematic representation of ClpXP. (B) ClpX domain structure and positions of sequence motifs important for ATP binding and hydrolysis (magenta), substrate binding (orange), and ClpP binding (yellow). (C) Left. Composite cryo-EM density of ClpX^ΔN^/ClpP complex. ClpP is yellow; ClpX^ΔN^ (class 3) is blue, green, or purple; and substrate is orange. Right. Slice through surface representation of the model showing substrate in the axial channel and the ClpP degradation channel. (D) Left. Cartoon representation of the class-4 ClpX^ΔN^ hexamer (subunit F removed) and its docking with a heptameric ClpP ring. Substrate in the ClpX channel is shown in space-filling representation as a poly-alanine chain; the inset shows substrate as a ball-and-stick model with associated density. The dashed line shows the 7-fold symmetry axis of ClpP. Right. View rotated by 90°. The dashed circle shows the position of the ClpP pore. (E) Cartoon representation of class-2 and class-4 ClpX^ΔN^ hexamers with substrate removed for clarity. Arrows point to the different positions of the seam interface.

ClpX subunits consist of a family specific N-terminal domain, which is dispensable for degradation of ssrA-tagged proteins, and large and small AAA+ domains, which contain sequence motifs that mediate ATP binding and hydrolysis, ClpP binding, and substrate recognition (Figure 1B; Baker and Sauer, 2012). Ring hexamers of ClpX bind the ssrA tag within an axial channel (Martin et al., 2008a). Following degron binding, ATP-fueled power strokes pull on and eventually unfold attached native domains (Kenniston et al., 2003). Single-chain ClpX^ΔN^ pseudohexamers, containing six ‘subunits’ linked by genetically encoded tethers, support ClpP degradation of ssrA-tagged substrates at rates similar to wild-type ClpX (Martin et al., 2005). Eliminating ATP hydrolysis in four or five subunits of single-chain pseudohexamers slows but does not prevent ClpP-mediated degradation, suggesting that ATP hydrolysis in any one of multiple subunits in the ClpX ring is sufficient to power unfolding and translocation (Martin et al., 2005). In optical-trapping experiments using single-chain ClpX and ClpP, the smallest observed translocation steps correspond to movement of five to eight amino acids of the substrate, and kinetic bursts of power strokes produce fast translocation steps two, three, or four-fold larger in terms of the number of residues translocated (Aubin-Tam et al., 2011; Maillard et al., 2011; Sen et al., 2013; Cordova et al., 2014; Olivares et al., 2017). Such bursts do not occur in repeating patterns, supporting probabilistic but coordinated ATP hydrolysis within the ClpX ring.

We describe here near atomic-resolution single-particle cryo-EM structures of single-chain ClpX pseudohexamers bound to ClpP and protein substrates. These structures show how asymmetric hexameric rings of ClpX dock with symmetric heptameric rings of ClpP and reveal how the pore-1, pore-2 and RKH loops of ClpX function in substrate binding. ClpX adopts a spiral conformation, with neighboring pore-1 loops interacting with every two residues of substrate in the axial channel, as observed in other AAA+ unfolding and remodeling machines (Puchades et al., 2017; de la Peña et al., 2018; Dong et al., 2019; Majumder et al., 2019; Ripstein et al., 2017; Gates et al., 2017; Zehr et al., 2017; Han et al., 2017; Su et al., 2017; Sun et al., 2017; Yu et al., 2018; White et al., 2018; Cooney et al. 2019; Rizo et al., 2019; Shin et al., 2019; Twomey et al., 2019). Based on these structural features, strictly sequential models of ATP hydrolysis and a power stroke that moves two residues of the substrate per translocation step have been proposed. As noted, however, ClpX does not need to operate in a strictly sequential manner and takes translocation steps substantially longer than two residues. Thus, an apparent incongruity exists between the structural and functional studies. We discuss this conflict and propose structure-based translocation models that reconcile how ClpX might use probabilistic ATP hydrolysis to take larger translocation steps of varying length.

## RESULTS

### Cryo-EM structures

For cryo-EM studies, we used epitope-tagged variants of *Escherichia coli* ClpP and a single-chain variant of *E. coli* ClpX^ΔN^ with an E185Q mutation to eliminate ATP hydrolysis without compromising nucleotide, ClpP, or substrate binding (Hersch et al., 2005; Martin et al., 2005). Single-chain ClpX^ΔN^ was used to ensure subunits of the pseudohexamer do not dissociate during sample preparation. This enzyme has been used previously for many biochemical and single-molecule studies (Martin et al., 2005; 2007; 2008a; 2008b; Aubin-Tam et al., 2011; Maillard et al., 2011; Glynn et al., 2012; Sen et al., 2013; Stinson et al., 2013; Cordova et al., 2014; Iosefson et al., 2105a; 2015b; Olivares et al., 2017; Rodriguez-Aliaga et al., 2016; Amor et al., 2019; Bell et al., 2018). We purified enzymes separately and incubated ClpX^ΔN^ (4 µM pseudohexamer), ClpP (2 µM 14-mer), and ATPγS (5 mM) for five min before vitrification. Experiments using ATP instead of ATPγS resulted in fewer ClpXP complexes. Imaging revealed complexes with ClpP bound to one or two ClpX^ΔN^ hexamers (Figure S1A-C). We analyzed the more abundant doubly capped complexes. As single-particle data processing with C_2_ symmetry produced maps with conformational heterogeneity, we used signal-subtraction methods to allow independent classification and refinement of ClpX^ΔN^ or ClpP density (Figures S1D-F). We calculated a D_7_ symmetric map for ClpP and parts of ClpX^ΔN^ making symmetric contacts, four different classes of symmetry-free ClpX^ΔN^ maps, and then extended each asymmetric ClpX^ΔN^ map to include one heptameric ClpP ring. Final structures have good stereochemistry with resolutions from 3.2 to 4.3 Å (Figure S2; Table S1). Substrates were observed in all ClpX^ΔN^ structures and probably represent bound endogenous peptides/proteins or partially denatured portions of ClpP or ClpX^ΔN^.

Figure 1C shows density for a composite ClpX^ΔN^/ClpP/substrate complex. The six subunits of ClpX^ΔN^ formed a shallow spiral (labeled ABCDEF from top to bottom) in all structural classes, which differed largely in substrate density or nucleotide state, and the hexameric ClpX^ΔN^ and heptameric ClpP rings were slightly offset (Figure 1D). The structure of the ClpX^ΔN^ hexamers in the class-1, class-3, and class-4 EM structures were very similar to each other (pair-wise Cα RMSDs 1.2-1.9 Å). The hexamer in the class-2 structure was generally similar (pair-wise Cα RMSDs 2.6 Å) but differed in the position of the ‘seam’, a dilated inter-subunit interface that occurs as a result of ring closure. This seam was located between subunits A and B in the class-2 structure and between subunits F and A in the class-1, class-3, and class-4 structures, (Figure 1E). As discussed below, this difference appears to be related to the identity of the nucleotide bound in subunits A or F.

### ClpX docking with ClpP

Our D_7_-symmetric map included ClpP_14_ and symmetric interface contacts with ClpX^ΔN^ (Figure 2A). In both heptameric ClpP rings, the N-terminal residues of each subunit formed a collar of β-hairpins creating a pore into the degradation chamber (Figures 2A-B, S3A). ClpP had essentially the same structure in the D_7_ and symmetry-free ClpXP maps. ClpX IGF loops, named for an Ile^268^-Gly^269^-Phe^270^ sequence, are critical for ClpP binding (Kim et al., 2001; Joshi et al., 2004; Martin et al., 2007; Amor et al., 2019) and were responsible for most contacts in our structures (Figure 2C). These interactions included packing of the Ile^268^, Phe^270^, and Val^274^ side chains into pockets at interfaces between ClpP subunits (Figure 2D). I268L, F270L, and V274A variants have severe ClpP binding defects (Amor et al., 2019). Asymmetric ClpX-ClpP docking relies on conformational adjustments in the N- and C-terminal residues of individual IGF loops, allowing the central portion of each IGF loop to contact the flat ClpP ring despite projecting from a spiral (Figure 2E). Changes in IGF-loop length decrease ClpP affinity and degradation activity (Amor et al., 2019), suggesting that these loops act as shock absorbers to maintain ClpP contacts during ClpXP machine function. In the symmetry-free maps, the unoccupied pocket in each ClpP heptamer was always located between the pockets bound by IGF loops from ClpX subunits E and F (Figure S3B; Movie S1).

**Figure 2.**
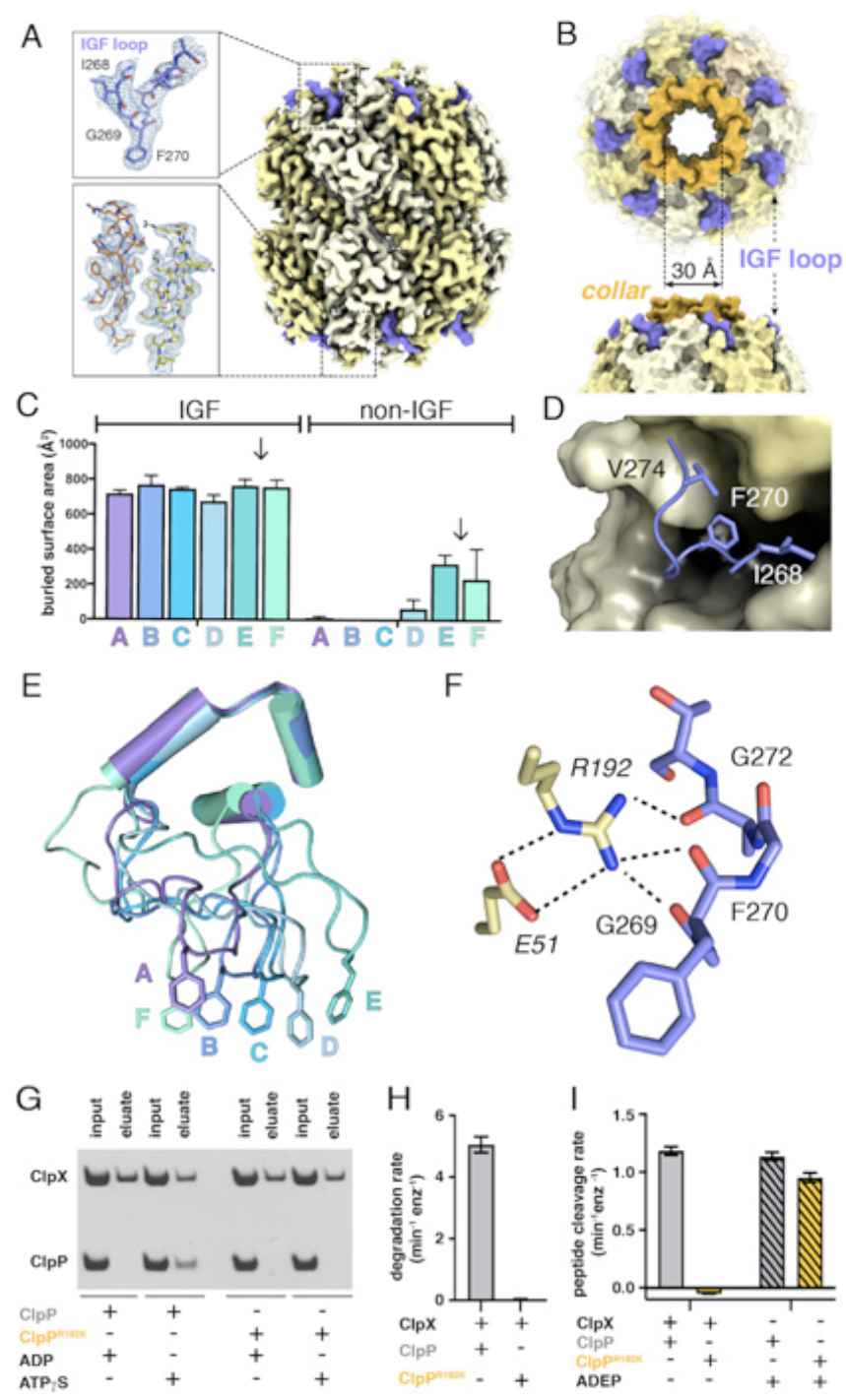
ClpP and symmetric IGF loop contacts. (A) Density for ClpP_14_ in the D_7_ map is shown in yellow. Density for part of the IGF loops of ClpX is shown in blue. The upper inset shows the density as mesh and the fitted model for residues 267-275 of one IGF loop; the lower inset shows density and the fitted model for resides 2-18 of two β-hairpins in the ClpP collar. (B) Top views of ClpP showing the axial collar (darker yellow) and pore. (C) IGF loops of ClpX make the majority of contacts with ClpP as assessed by buried surface area. Each bar represents a different ClpX^ΔN^ subunit in the spiral and values are means ± 1 SD for the four classes. The arrows between subunits E and F mark the position of the unoccupied ClpP cleft. (D) The side chains of IGF residues Ile^268^ (I268) Phe^270^ (F270), and Val^274^ (V274) pack into hydrophobic ClpP clefts. (E) After alignment of the large AAA+ cores of the six subunits in the class-4 ClpX^ΔN^ hexamer, the IGF loops adopt a variety of conformations, helping mediate ClpP binding despite the symmetry mismatch. The side chains of Phe^270^ are shown in stick representation with the IGF loop (residues 263-283) and three flanking helices (residues 254-262 and 284-298) shown in cartoon representation. (F) The side chain of Arg^192^ (R192) in ClpP makes hydrogen bonds with carbonyl oxygens in the IGF loop of ClpX and also forms a salt bridge with Glu^51^ (E51) in a neighboring ClpP subunit. (G) ClpP but not ^R192K^ClpP binds ClpX^ΔN^ in pull-down assays performed in the presence of ATPγS. As a negative control, neither ClpP variant binds ClpX^ΔN^ in the presence of ADP (Joshi et al., 2004). (H) ClpX^ΔN^ supports degradation of ^cp7^GFP-ssrA by ClpP but not by ^R192K^ClpP. (I) ClpX^ΔN^ supports degradation of a decapeptide by ClpP but not by ^R192K^ClpP. ADEP-2B activates decapeptide cleavage by both ClpP and ^R192K^ClpP. Error bars represent means (n=3) ± 1 SD.

In our structures, the side chain of Arg^192^ in ClpP appeared to hydrogen bond to the backbone of the IGF loop (Figure 2F). Unlike wild-type ClpP, ^R192K^ClpP neither bound ClpX^ΔN^ in pull-down assays (Figure 2G) nor degraded protein substrate in the presence of ClpX^ΔN^ (Figure 2H). Small-molecule acyldepsipeptides (ADEPs) bind in the same ClpP pockets as the ClpX IGF loops and kill bacteria by opening the ClpP pore to facilitate rogue degradation of unstructured proteins (Brötz-Oesterhelt et al., 2005; Kirstein et al., 2009; Li et al., 2010; Lee et al., 2010b). ADEPs stimulated R192K and wild-type ClpP decapeptide cleavage to similar extents (Figure 2I), establishing that ^R192K^ClpP has normal peptidase activity. Thus, Arg^192^ is critical for ClpX binding but not for ADEP binding or ClpP pore opening. Modifying ADEPs to interact with Arg^192^ may increase affinity for ClpP and improve their efficacy as antibiotics.

Crystal structures show that ADEP binding to ClpP results both in opening of the axial pore and assembly of its N-terminal sequences into a collar of β-hairpins (Li et al., 2010; Lee et al., 2010b). Based on a recent cryo-EM structure of *Listeria monocytogenes* ClpXP, it was concluded that ClpX binding does not induce the same widening of the ClpP pore as does ADEP binding (Gatsogiannis et al., 2019). By contrast, our results support the opposite conclusion, as the diameter and overall structure of the ClpP pore in our *E. coli* ClpXP structures were extremely similar to the crystal structure of ADEP-activated *E. coli* ClpP (Li et al., 2010), with an overall RMSDs of 0.8 Å for all Cα’s in a single ClpP ring and 0.6 Å for all Cα’s of the seven N-terminal β-hairpins and adjacent α-helices that define the pore diameter.

### Substrate binding

All ClpX^ΔN^ maps contained substrate density within the axial channel, which we generally modeled as an extended poly-alanine chain (Figure 1D), although additional side-chain density at residue 3 of the substrate was modeled as arginine in the class-1 structure and histidine in the class-3 structure (Figure S4A). Substrate was built with the N terminus facing ClpP in the class-2 and class-4 structures and in the opposite orientation in the class-1 and class-4 structures. ClpXP can translocate substrates in either the N-to-C or C-to-N direction (Olivares et al., 2017). In all structures, the top of the ClpX channel was most constricted by tight packing between ClpX side chains and substrate (Figures 3A-B), providing a structural basis for experiments showing that interactions with substrate near the top of the ClpX channel are most important for unfolding grip (Bell et al., 2019). The class-1 and class-3 structures also had density corresponding to a roughly globular native domain above the channel (Figures 1C, S4B).

**Figure 3.**
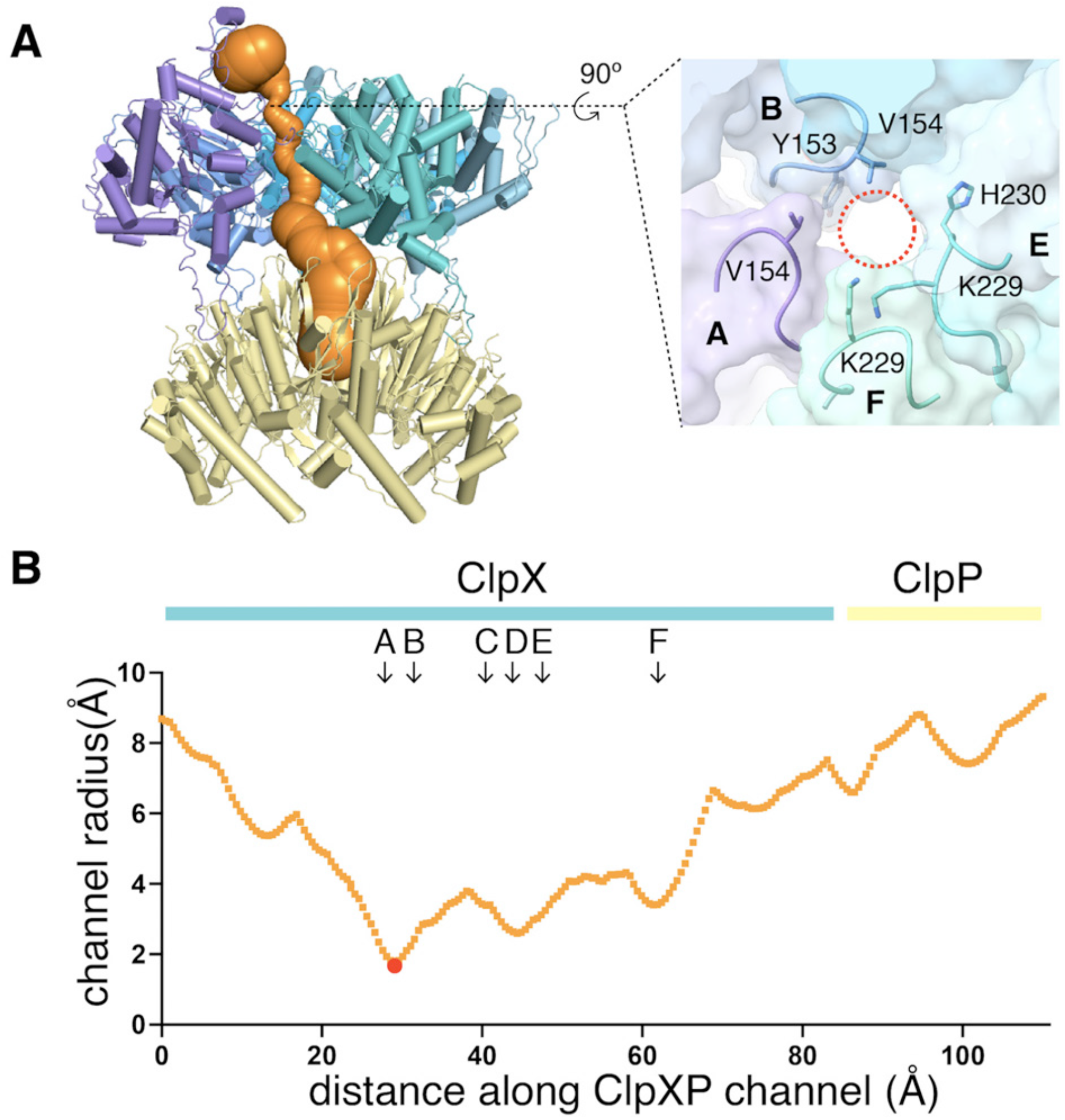
Dimensions of axial channel. (A) The channel though ClpX and into ClpP (class-1) is most constricted near the top of the ClpX^ΔN^ hexamer. This panel was made using *Caver* (Pavelka et al., 2016). The inset shows that the constriction involves the pore-1 loops of subunits A and B and the RKH loops of subunits E and F. (B) Plot of channel radius as a function of channel length. Arrows mark positions of pore-1 loops. Red dot marks the bottleneck of this channel.

In our structures, the pore-1, pore-2, and RKH loops of ClpX contacted substrate in accord with genetic and biochemical studies (Siddiqui et al., 2004; Farrell et al., 2007; Martin et al., 2008a; 2008b; Iosefson et al., 2015a; 2015b; Rodriguez-Aliaga et al., 2016). For example, the pore-1 loops of subunits B, C, D, and E interacted with substrate in the channels of all classes, whereas these loops from subunits A and F did so only in some classes (Figures 4A-C, S4C-D; Movie S2). For each engaged pore-1 loop, the side chains of Tyr^153^ and Val^154^ packed between β-carbons spaced two-residues apart on opposite sides of the extended substrate (Figures 4A-C, S4C-D; Movie S2). Depending on the class, from three to five pore-2 loops also contacted substrate, with Arg^200^ making most of these interactions (Figures 4A, 4B, 4D, S4C-D; Movie S2).

**Figure 4.**
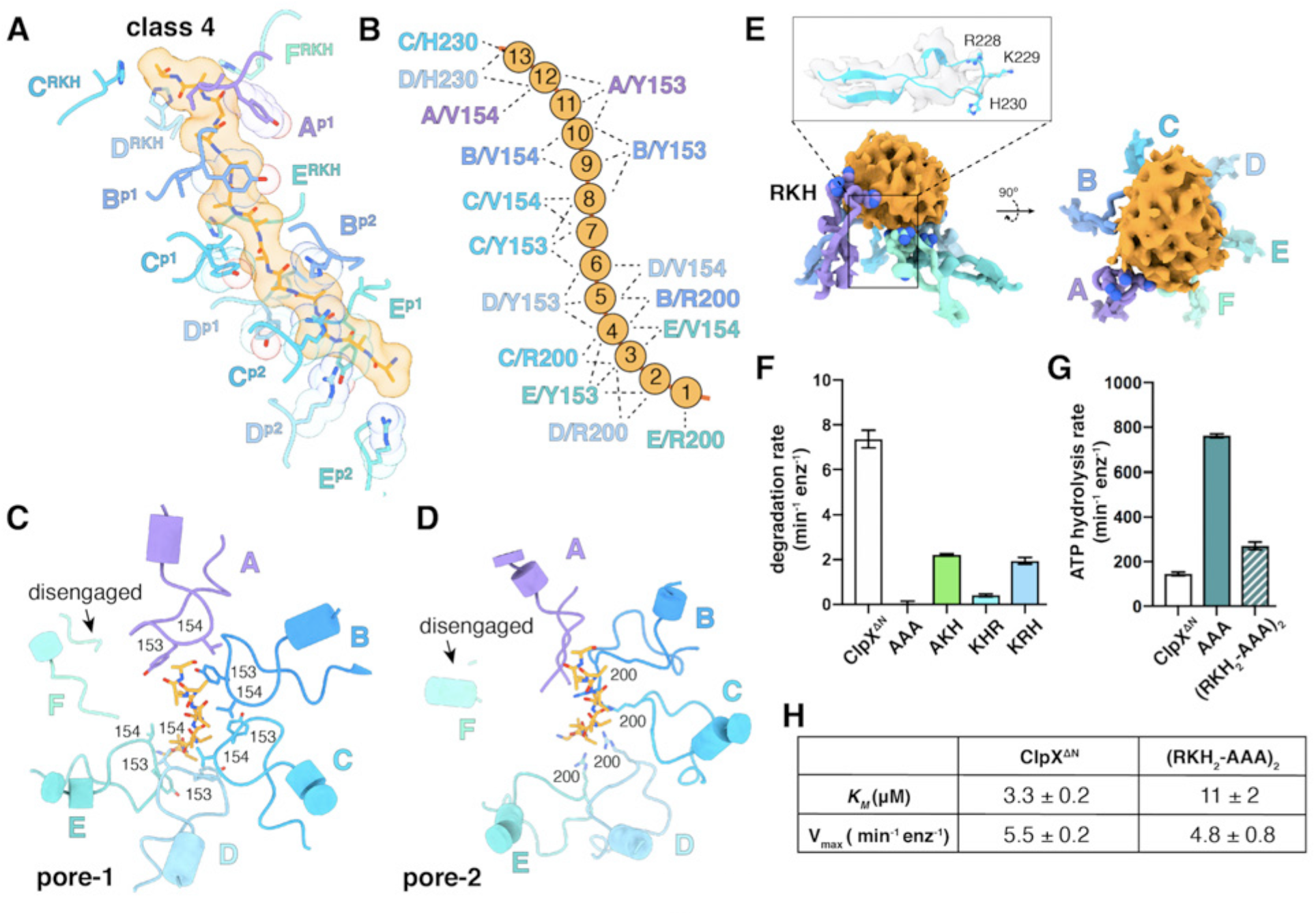
Substrate contacts. (A) Interactions between substrate (orange, stick and surface representation) and ClpX^ΔN^ loops (blue and purple, stick and transparent space-filling representation) in the class-4 structure. Capital letters indicate subunits; superscripts indicate RKH loops, pore-1 (p1) loops, or pore-2 (p2) loops. (B) Scheme of interactions shown in panel (A). Dashed lines represent distances of 6.5 Å or less between the Cβ atoms of substrate alanines and the Cβ atoms of Y153/V154 (p1), or the Cε atom of R200 (p2), or the Cγ atom of H230 (RKH). (C) Top view of interactions of pore-1 loops with substrate (class 4). (D) Top view of interactions of pore-2 loops with substrate (class 4). (E) Interaction of RKH loops in the class-3 structure with the globular portion of the substrate above the channel. Inset – representative RKH-loop density (class 4, subunit C) and positions of R228, K229, and H230. (F) Mutation of RKH motifs in each subunit of a ClpX^ΔN^ hexamer inhibits degradation of Arc-st11-ssrA. (G) Effects of RKH mutations on ATP hydrolysis. (H) Mutating two RKH motifs in a single chain pseudohexamer to AAA increases *K*_M_ for steady-state degradation of ^CP7^GFP-ssrA.

The RKH loop, which has not been visualized in previous structures and is unique in ClpX-family enzymes (Baker and Sauer, 2012), consisted of an antiparallel β-ribbon stem and a short helix that includes part of the conserved Arg^228^-Lys^229^-His^230^ motif (Figure 4E). When contacting substrate above the pore, the RKH loops mimicked adjustable structural jacks supporting a house during foundation repair (Figure 4E). The RKH loops alter substrate specificity (Farrell et al., 2007; Martin et al., 2008a), but whether they are critical determinants of substrate binding is unknown. To test this possibility, we constructed and assayed RKH-loop mutants. The most severe mutant (RKH◊AAA) did not support ClpP degradation of an ssrA-tagged substrate and hydrolyzed ATP ∼5-fold faster than the parent (Figures 4F-G). Changing RKH to AKH, KHR, or KRH slowed degradation (Figure 4F). A variant [RKH_2_-AAA]_2_ pseudohexamer with four wild-type RKH loops and two AAA mutations had a ∼3-fold higher *K*_M_ for substrate degradation than the parent and hydrolyzed ATP slightly faster (Figures 4G-H). Thus, the RKH loops play important roles in substrate recognition and in regulating rates of ATP hydrolysis.

### Nucleotide binding and motor conformations

We observed density for five ATPγS molecules in each ClpX^ΔN^ hexamer (Figure 5A; Table S2). In the sixth site, bound nucleotide fit best as ADP in subunit A (class 2) or subunit F (classes 1, 3, and 4), although with slightly less convincing density than for ATPγS in other subunits (Fig. 5B; Table S2), raising the possibility of averaging of multiple nucleotide-binding states. These ‘ADP-bound’ subunits had poorly structured channel loops, made fewer contacts with substrate and with neighboring subunits, and were offset from the axis of the hexamer by ∼2.5 Å (Figures 5C-5E; Table S3).

**Figure 5.**
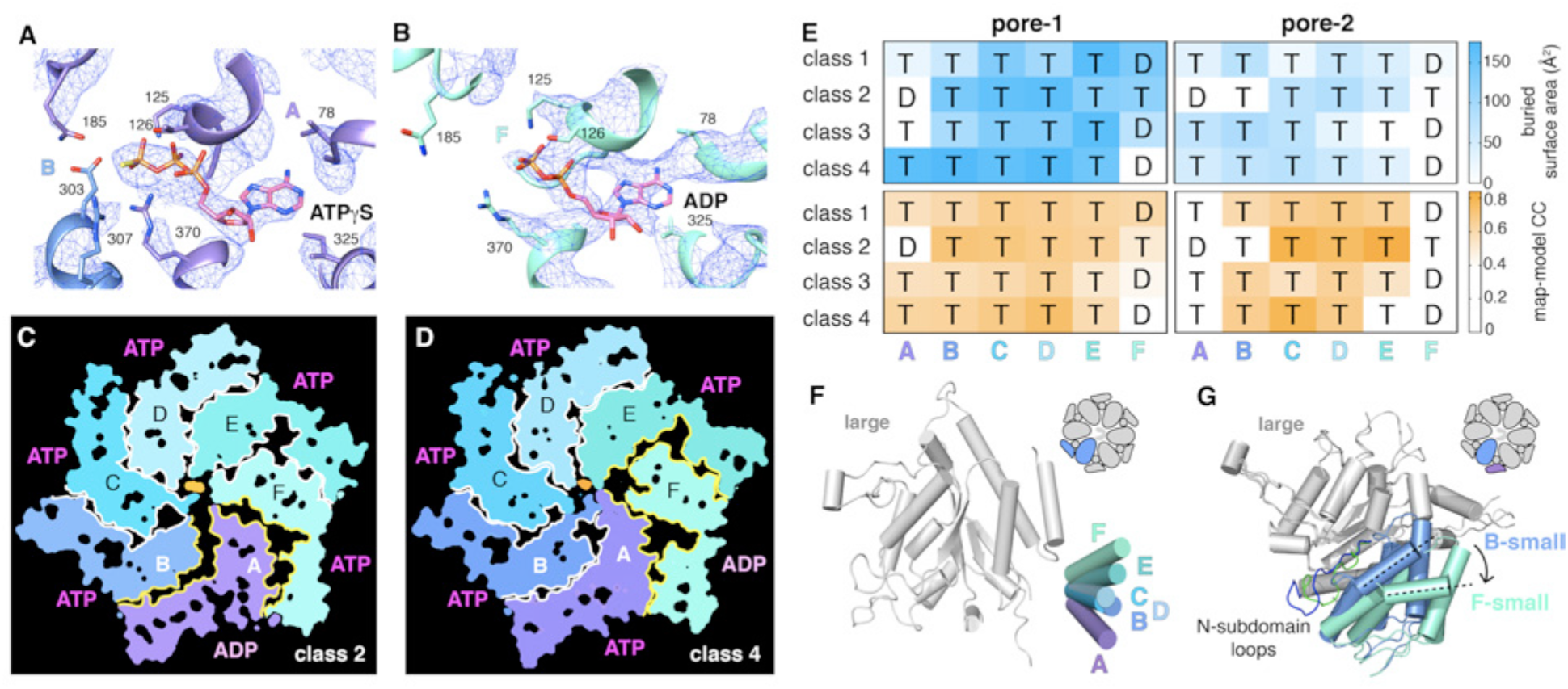
Nucleotide binding and subunit interactions in ClpX^ΔN^ hexamers. (A) Density for ATPγS in subunit A of the class-4 hexamer. The positions of ATP-binding/hydrolysis motifs are shown in cartoon and stick representation. (B) Weaker nucleotide density built as ADP in subunit F of the class-4 hexamer. (C, D) Slices through the density maps of class-2 and class-4 hexamers show that the ADP-bound subunit (A or F, respectively) makes fewer neighbor contacts than ATP-bound subunits (yellow). (E) Contacts between substrate and pore-1 loops (left) or pore-2 loops (right) were characterized by buried surface area (top) or model-to-map correlation (bottom). T represents an ATP-bound subunit; D represents an ADP-bound subunit. (F) Superimposition of the large AAA+ domains of six subunits from a hexamer (class 4) showing variation in the angle between small and large domains of the same subunit. One of the large AAA+ domains is light grey, and the small AAA+ domains are in different shades of purple, blue and green. (G) Superimposition of two large AAA+ domains from a class-4 hexamer, illustrating different types of packing against the neighboring small AAA+ domain. The large domains of subunits A and C are light grey; the small domains of subunit F and B are cornflower blue and mint green. N sub-domain loops are colored green (subunit F) and blue (subunit B). The dashed lines and arrow show an ∼30° rotation at the interface. The F/A interface is the seam.

Side-chain contacts with nucleotide included ClpX residues Val^78^-Ile^79^ (box-II), Lys^125^-Thr^126^-Leu^127^ (Walker A), Asp^184^-Gln^185^ (Walker B with E185Q mutation), Arg^307^ (arginine finger), and Arg^370^ (sensor-II) (Figures 5A, 5B, S5A). V78A/I79A, E185Q, and R370K mutations eliminate ATP hydrolysis or weaken nucleotide binding (Joshi et al., 2004; Hersch et al., 2005; Martin et al., 2005; Stinson et al., 2013). We found that K125M, T126A, L127A, D184A, and R307A mutations also severely inhibited ATP hydrolysis (Figure S5B). ClpX does not have a traditional sensor-I residue, which in many AAA+ enzymes positions a water molecule for hydrolysis of ATP bound to the same subunit (Erzberger and Berger, 2006). For ATPγS bound in subunits A-E of our structures, the side chain of Glu^303^ was close to the γ-thiophosphate, at a sensor-I-like position (Figure 5A). We found that an E303A mutation severely inhibited ATP hydrolysis, whereas E303Q was partially defective (Figure S5B-C). Based on these results, we refer to Glu^303^ as a sensor-I* element. As Glu^303^ is in the same helix as Arg^307^, the arginine-finger residue, these residues could coordinate structurally to activate ATP hydrolysis in a neighboring subunit.

A short hinge connects the large and small AAA+ domain of each ClpX^ΔN^ subunit. In all of our structures, the conformations of the small domains were very similar to each other, as were those of the large domains after removing an N-terminal subdomain (residues 65-114) and the pore-1, pore-2, RKH, and IGF loops (Figure S6). The relative orientations of the large and small AAA+ domains were similar across all four structural classes for subunits at equivalent spiral positions, but conformational changes in the hinge resulted in different orientations or the large and small domains at many positions in the spiral for each hexamer (Figure 5F). Changes in hinge length or deletion of one hinge largely eliminate ClpX function (Glynn et al., 2012; Bell et al., 2018). Thus, the conformational changes associated with a power stroke likely arise from changes in hinge conformations, whereas movements of different large-domain loops or the N-subdomain mediate ring closure and asymmetric contacts with ClpP and substrate.

ATP hydrolysis requires proper positioning of the box-II, Walker-A, and Walker-B elements in the large AAA+ domain of one ClpX subunit, the sensor-II arginine from the small AAA+ domain of the same subunit, and the arginine-finger/sensor-I* element from the large domain of the clockwise subunit (viewed from the top of the ring). These structural features depend on how the small AAA+ domain of each subunit packs against the large AAA+ domain of its clockwise neighbor, which was similar for units A/B, B/C, C/D, D/E, and E/F, suggesting that the associated ATP-binding sites are hydrolytically active. By contrast, the structure of the FA interface was different as a consequence of changes in rotation of the N-subdomain in subunit A relative to the rest of the large AAA+ domain and changes in a loop that contains the sensor-II arginine in subunit F (Figure 5G). These changes resulted in disengagement of the Arg-finger and sensor-I* side chains in subunit A from the nucleotide bound to subunit F (Figure 5B; Table S4). Thus, in each of our structures, the nucleotide-binding site in subunit F appears to be catalytically inactive.

## DISCUSSION

### ClpX interactions with ClpP and substrates

Our cryo-EM structures provide snapshots of ClpX binding to ClpP, protein substrates, and nucleotides. In each of our structures, the six subunits of the ClpX ring hexamer are arranged in a shallow spiral. Slightly altered orientations of the large and small AAA+ domains in each ClpX subunit allow the hexameric ring to remain topologically closed with each large domain contacting the small domain of one neighbor. These structural results are consistent with biochemical experiments that show that ClpX is fully functional when all of its subunit-subunit interfaces are covalently crosslinked (Glynn et al., 2012) and support a model in which the architectural changes in the spiral that drive a power stroke result from changes in the conformations of the hinges connecting the large and small AAA+ domains of each subunit. Our structures show that relatively flexible interactions between IGF loops of ClpX and binding pockets on ClpP heptamers allow docking of these symmetry-mismatched partners. Although IGF-ClpP contacts are highly dynamic in solution (Amor et al., 2016), they were well defined in our structures and revealed essential interactions.

Protein substrates were observed in the axial channel of ClpX in all structures and above the channel in two structures. The ClpX pore-1 and pore-2 loops were responsible for most substrate contacts in the channel, with a periodicity of two substrate residues per ClpX subunit. The RKH loops of ClpX, which we found play critical roles in substrate recognition and control of ATP-hydrolysis rates, also contacted substrate near the top and above the channel. Substrate contacts near the top of the channel are most important in determining substrate grip during unfolding (Bell et al., 2019), and substrate-ClpX contacts were tightest in this region in our structures. Finally, we observed ATPγS bound to five of the six ClpX subunits with ADP apparently bound to the sixth subunit, and determined how ClpX functions without a traditional sensor-I residue.

Our cryo-EM structures contrast markedly with crystal structures of ClpX^ΔN^ pseudohexamers, which did not form a spiral, bound only four nucleotides, and had conformations that now seem incompatible with ClpP and/or substrate binding (Glynn et al., 2009; Stinson et al., 2013).

### Proteolytic motors from different AAA+ clades have similar structures

AAA+ protease motors belong to either the classic or HCLR clades (Erzberger and Berger, 2006). For HCLR-clade members ClpX and Lon, the spiral hexamer architectures and pore-1-loop interactions with substrate in the axial channel are similar to those for the classic-clade YME1/FtsH and the proteasomal Rpt_1-6_/PAN motors (Puchades et al., 2017; de la Peña et al., 2018; Dong et al., 2019; Majumder et al., 2019; Shin et al, 2019). Thus, from a structural perspective, it is reasonable to suggest that motors from different clades may operate by a common fundamental mechanism.

A sequential translocation model has been proposed for Yme1 and the Rpt_1-6_ ring of the 26S proteasome, based on placing distinct cryo-EM structures with different nucleotide and substrate-engagement states in a defined kinetic pathway, (Puchades et al., 2017; de la Peña et al., 2018; Dong et al., 2019). In this model, ATP hydrolysis in the fifth spiral subunit (subunit E in ClpX) drives a power stroke that moves each subunit from its previous location to a position offset by one subunit in the clockwise direction, generating a two-residue translocation step (Figure 6A). We refer to this translocation model as SC/2R (<u>S</u>equential <u>C</u>lockwise/<u>2</u>-<u>R</u>esidue Step). Based on similar hexamer architectures and/or substrate interactions, a SC/2R translocation model has also been proposed for Lon, for PAN/20S, and for the AAA+ protein-remodeling machines ClpB/Hsp104, CDC48/p97, NSF, and Vps4 (Gates et al., 2017; Han et al., 2017; Ripstein et al., 2017; Su et al., 2017; Sun et al., 2017; Zehr et al., 2017; Yu et al., 2018; White et al., 2018; Cooney et al. 2019; Majumder et al., 2019; Rizo et al., 2019; Twomey et al., 2019; Shin et al., 2019).

**Figure 6.**
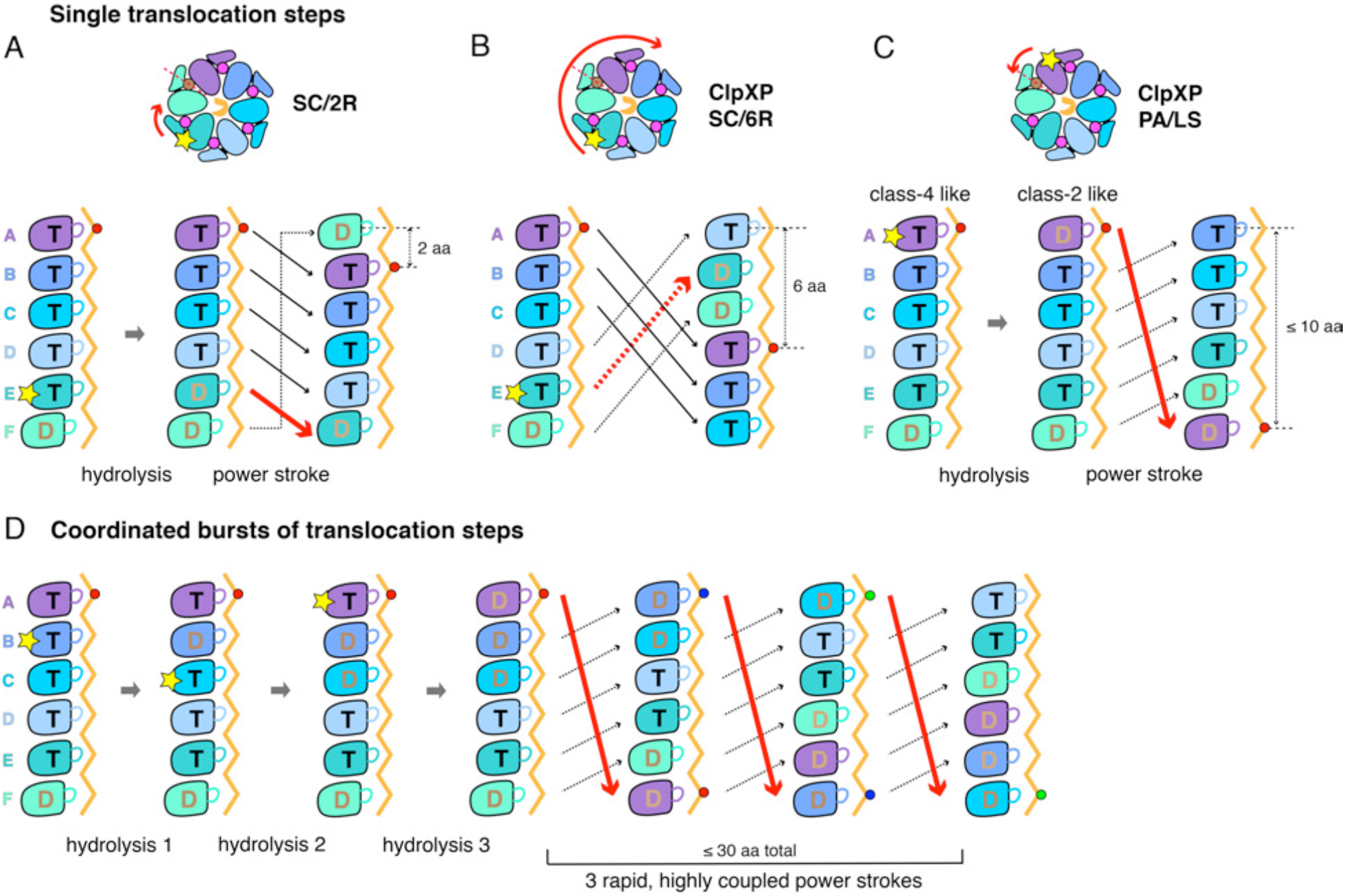
Single-step and burst translocation models. In all panels, T represents ATP-bound subunits, and D represents subunits containing ADP and possibly inorganic phosphate. (A) SC/2R translocation model proposed for the Yme1 protease and 26S proteasome (Puchades et al., 2017; de la Peña et al., 2018; Dong et al., 2019). Only subunit E hydrolyzes ATP (depicted by a star). Hydrolysis and/or product release results in a two-residue translocation step, in which the top five subunits in the spiral move down one position in the clockwise direction and the bottom subunit moves up to the top position. (B) SC/6R translocation model. During a six-residue translocation step, subunits A, B, and C each move down in the spiral to positions D, E, and F, respectively, dragging substrate with them; at the same time, subunits D, E and F each move up to positions A, B, and C. Subunit displacement is in the clockwise direction. (C) PA/LS model in which ATP hydrolysis in subunit A results in a single translocation step of up to 2.5 nm in length as a consequence of anti-clockwise movement of this subunit to position F at the bottom of the spiral. At the same time, subunits BCDEF move up to positions ABCDE. (D) One variation of a PA/LS model resulting in a burst of three long translocation steps. Initial probabilistic ATP hydrolysis in subunits D and then B creates strain in the spiral, which is released in burst of fast steps upon ATP hydrolysis in subunit A. In the general PA/LS model, the initial ATP-hydrolysis event can occur with different probabilities in subunits A-E.

### Discrepancies between ClpX function and SC/2R-model predictions

The ClpX structures reported here resemble those of Yme1 and Rpt_1-6_ in the spiral architecture of the hexamer, the positions of subunits that contain a nucleoside triphosphate, the interaction of successive pore-1 loops with two-residue segments of the substrate, and the patterns of substrate engaged and disengaged pore-1 and pore-2 loops in the ring (Puchades et al., 2017; de la Peña et al., 2018; Dong et al., 2019). Thus, ClpXP might also be expected to operate by an SC/2R mechanism, but multiple experimental observations suggest otherwise.

The first issue involves the length of the smallest, fundamental translocation steps. Structures show that axial-channel binding in the AAA+ ring enforces an extended polypeptide conformation in which two residues of the substrate span ∼0.6 nm, and the SC/2R model predicts corresponding step sizes in terms of two residues translocated or ∼0.6 nm traversed. By contrast, the basic translocation step of ClpXP measured in optical-trapping experiments is ∼1 nm (Aubin-Tam et al., 2011; Maillard et al., 2011; Sen et al., 2013; Cordova et al., 2014; Iosefson et al., 2015b; Rodriguez-Aliaga et al., 2016; Olivares et al., 2017). Although ∼1 nm is larger than ∼0.6 nm, this difference, by itself, is not a compelling argument against the SC/2R model. In the optical trap, however, step size represents the average distance that unfolded polypeptide outside of the axial channel moves between successive translocation steps, which can be converted into amino-acid residues using the wormlike-chain model (Bustamante et al., 1994). Because the unfolded substrate outside the channel is in a partially compact conformation at the forces used in these experiments, ∼1 nm corresponds to a translocation step of 5-8 residues, which appears inconsistent with the two-residue step predicted by the SC/2R model.

Because the sensitivity of optical trapping precludes direct identification and quantification of translocation steps as short as two residues, it could be argued that the single 5-8 residue steps observed for ClpXP actually consist of three or four unresolved SC/2R sub-steps. Consideration of kinetics makes this possibility unlikely, however. For example, the time from the beginning to the end of each translocation step is less than 0.1 s in the optical trap (12-15). Under similar conditions, the steady-state rate of ClpX^ΔN^/ClpP ATP hydrolysis is 3.6 ± 0.1 s^−1^ (Fig. S7), corresponding to a time constant of 0.28 s. In the SC/2R model, each sub-step would require an independent ATP hydrolysis event, and ∼1 s would be required to take three or four sub-steps. This kinetic problem becomes worse for ClpXP translocation bursts that move 20 to 32 residues in less than 0.1 s. Thus, ClpXP translocation steps occur ∼10 to ∼40-fold faster than predicted by the SC/2R mechanism and the experimentally determined rate of ATP hydrolysis.

Another caveat might be that ClpX step sizes could be longer under tension in optical-trapping experiments, or that ClpXP uses a different translocation mechanism under these conditions. We consider both possibilities unlikely as the distribution of translocation step lengths shows little dependence on trap force (Aubin-Tam et al., 2011; Maillard et al., 2011; Sen et al., 2013; Cordova et al., 2014; Olivares et al., 2017).

A separate concern involves the SC/2R prediction that only subunit E in the ClpX spiral hydrolyzes ATP during normal translocation, and thus that single-chain variants containing subunits that cannot hydrolyze ATP should stall when an ATPase inactive subunit moves into the hydrolysis position. However, ClpX^ΔN^ hexamers with just two ATPase-active subunits support ClpP degradation at ∼30% of the wild-type rate and with the same thermodynamic efficiency as wild-type ClpXP (Martin et al., 2005). Thermal motions might move an ATPase-inactive subunit out of the hydrolysis position in the ClpX spiral and an active subunit into it, allowing continued function. This possibility seems unlikely, as the variant with just two ATPase active subunits translocates substrates directionally against force in optical-trap experiments (Cordova et al., 2014). Thus, at some level, ClpX translocation must operate probabilistically to avoid stalling in cases where only some of its subunits are hydrolytically active.

### Structure-based translocation models for ClpXP

To account for the experimentally determined translocation properties of ClpXP, any model needs to explain: (1) how structural changes in the spiral result in a fundamental translocation step of 5-8 residues despite the observed two-residue periodicity of substrate contacts; (2) how kinetic bursts could generate very fast translocation of 20-32 residues without requiring multiple ADP-dissociation and ATP-rebinding events; and (3) how the motor functions efficiently without strict requirements for ATP hydrolysis in any particular subunit in the spiral.

Modifications of the SC/2R model could, in principle, address some of these issues. For example, upon ATP hydrolysis in subunit E, a clockwise movement a three-subunit shift could generate a six-residue power stroke in a SC/6R model (Figure 6B). Potential issues with this mechanism include that substrate contacts with subunits D, E, and possibly F would need to break during each translocation step; the structural features that would drive a three-subunit rearrangement are unclear; and this SC/6R model doesn’t account for kinetic bursts resulting in translocation steps longer than six residues. By altering this clockwise three-subunit shift mechanism to allow probabilistic hydrolysis, long translocation bursts can be explained. Specifically, assume that multiple subunits in the spiral have some probability of hydrolyzing ATP, with the stipulation that hydrolysis at subunits other than E causes strain that is only released in a burst of six-residue power strokes when ATP in subunit E is eventually hydrolyzed or sufficient strain in the spiral accumulates (<u>P</u>robabilistic <u>C</u>lockwise/<u>6</u>-<u>R</u>eside Step or PC/6R model). Five of the six nucleotide-binding sites in our ClpX structures have similar geometries and thus each of these sites could plausibly hydrolyze ATP. Probabilistic models do not preclude neighboring subunits in the ring from firing sequentially, but rather exclude any mechanistic requirement for sequential action, and are fully compatible with communication between ClpX subunits and cooperative ATP hydrolysis (Hersch et al., 2005; Martin et al., 2005).

Examination of our ClpXP structures reveals an alternative mechanism by which a fundamental translocation step of 5-8 residues might occur. Specifically, a power stroke that moves the top subunit in the ClpX spiral to the bottom position in the anti-clockwise direction could generate a movement of up to 2.5 nm through the axial channel. A conformational change of this type is captured in a morph in which subunit A in class 2 is aligned with subunit F in class 4 (Movie S3). Figure 6C shows the corresponding translocation model in which probabilistic ATP hydrolysis in subunit A results in a single long step (<u>P</u>robabilistic <u>A</u>nti-clockwise <u>L</u>ong-<u>S</u>tep or PA/LS). The long-step nomenclature implies that step size need not be fixed in terms of residues, and could depend on conformational variability in the substrate sequence. A similar anticlockwise A-to-F subunit transition has been proposed for the Vps4 unfoldase (Su et al., 2017). Probabilistic ATP hydrolysis at multiple subunits in the spiral could then account for a kinetic burst of translocation steps, obviating the need to release ADP and rebind ATP after each step. Figure 6D shows one variation of this model. In any PA/LS model, subunit A would need to bind substrate and drag it to the bottom of the spiral, at least transiently breaking contacts with other subunits. Whether these contacts would be physically broken or simply released during the power stroke is currently unclear, as not all conformational states in the ClpX reaction cycle are likely to be known at this point. Crystal structures of ClpX reveal rotations between the large and small AAA+ domains so large that nucleotide cannot bind some subunits (Glynn et al., 2009; Stinson et al., 2013). Although subunits of this type are not present in the current cryo-EM structures, they could be representative of transient functional conformations, as crosslinks engineered to trap these hexamer conformations form in solution and prevent ClpXP degradation but not ATP hydrolysis (Stinson et al., 2013; 2015).

Additional observations indicate that functionally relevant conformations remain to be discovered. For example, ClpXP degrades disulfide-bonded and knotted proteins in reactions that require simultaneous translocation of two or more polypeptides (Burton et al., 2001; Bolon et al., 2004; San Martín et al., 2017; Sivertsson et al., 2019). In our cryo-EM structures, a single polypeptide strand fills the axial channel, and thus structures must exist in which the channel expands to accommodate multiple strands during translocation.

### Other AAA+ unfolding/remodeling machines and SC/2R mechanisms

ClpAP takes ∼1 nm and ∼2 nm residue translocation steps (Olivares et al., 2014; 2017), suggesting that it can also operate by a non-SC/2R mechanism, but we are unaware of experiments that establish the translocation step size for other AAA+ protein unfolding or remodeling enzymes. Like ClpX, some of these enzymes do not stall if an ATPase-inactive subunit occupies the hydrolysis position. In the Rpt_1-6_ hexamer of the yeast 26S proteasome, for example, ATPase-defective Rpt_3_, Rpt_4_, or Rpt_6_ subunits cause virtually complete loss of degradation activity, whereas ATPase-defective Rpt_1_, Rpt_2_, and Rpt_5_ subunits do not (Beckwith et al., 2013). In the alternating Yta10/Yta12 hetero-hexamer (a Yme1 homolog), a Walker-B mutation in Yta12 prevents ATP hydrolysis in Yta10, as expected for tight subunit-subunit coupling in a strictly sequential mechanism, but a Walker-B mutation in Yta10 does not prevent ATP hydrolysis in Yta12 (Augustin et al., 2009). Given the strong similarities between the structures and substrate interactions of a large number of AAA+ unfolding and remodeling machines and the fact that the SC/2R mechanism does not account for ClpX and ClpA experimental results, it will be important to continue to use structural and functional studies to investigate the molecular mechanisms by which these AAA+ machines function.

## METHODS

### Protein expression and purification

The single-chain ClpX variant used for microscopy contained six copies of *E. coli* E185Q ClpX^ΔN^ (residues 62-424), with neighboring units connected by six-residue linkers of variable composition, and a C-terminal TEV cleavage site and His_6_ tag. The ClpP variant consisted of full-length *E. coli* ClpP followed by a TEV cleavage site and His_6_ tag. Both proteins were expressed separately in *E. coli* strain ER2566 and purified as described (Stinson et al., 2013). After purification, TEV protease was used to remove the His_6_ tags. TEV protease and uncleaved proteins were removed by Ni^2+^-NTA affinity chromatography, and purified proteins were flash frozen in storage buffer (20 mM HEPES, pH 7.5, 300 mM KCl, 0.5 mM EDTA, 10% glycerol) and stored at –80 °C.

### Sample preparation, data acquisition, and image processing

To assemble complexes, the ClpX^ΔN^ pseudohexamer and ClpP_14_ were diluted into EM buffer (20 mM HEPES, pH 7.5, 100 mM KCl, 25 mM MgCl_2_, 5 mM ATPγS) to final concentrations of 4 µM and 1.8 µM, respectively. After 5 min at 25 °C, 3 µL of the mixture was applied to glow discharged R1.2/1.3 400 mesh grids (Quantifoil). Grids were blotted with filter paper 494 (VWR) and plunged into liquid ethane using a Cryoplunge-3 system (Gatan). Electron micrographs were collected using a Talos Arctica with a Gatan K2-Summit direct electron detector in super-resolution mode. High-resolution movies were recorded at a magnification of 36000X (0.58 Å pixel size). Each movie was composed of fifty frames (200 ms per frame) and a total dose of ∼58 e^−^/Å^2^ per movie. The final dataset consisted of 3657 movies recorded in two separate sessions. Frames in each movie were 2X binned, aligned, gain-corrected, and dose-weighted using Motioncor2 (Zheng et al., 2017), to generate a single micrograph. The contrast transfer function (CTF) was estimated using CTFFIND4 (Rohou and Grigorieff, 2015). Unless noted, Relion 2.0 (Kimanius et al., 2016) was used for 2D/3D classification and refinement.

We first attempted to construct a density map of doubly capped ClpX-ClpP-ClpX using standard protocols. 1.4 million doubly capped particles were automatically picked and filtered by 2D classification. 443,717 “good” particles were selected for 3D map reconstruction. To generate an initial model for 3D refinement, the crystal structures of a ClpX^ΔN^ hexamer (PDB 3HWS; Glynn et al., 2009) and a ClpP tetradecamer (PDB code 3MT6; Li et al., 2010) were merged in PyMOL and low-pass filtered to 40 Å. 3D refinement with C_2_ symmetry yielded a map with a resolution of ∼4 Å, but the quality of this map was poor and interpretation of secondary structure elements was impossible. Using C_1_ or C_7_ symmetry did not improve the map.

To minimize problems caused by the symmetry mismatch between ClpX and ClpP, we treated ClpX and ClpP separately before the last step of refinement. For the ClpP reconstruction, we applied a soft circular mask, including ClpP and the tips of the ClpX IGF loops, to the 3D reference and 2D images, respectively. Because we observed predominantly side views of ClpXP (perpendicular to the axial channel; Figure S1A-C), this simple masking of the particle images allowed us to remove most of the ClpX density and to focus particle alignment on ClpP. Starting from a low-pass filtered ClpP structure as an initial reference, 3D refinement with D_7_ symmetry yielded a 3.2-Å resolution map with clear secondary structure and side-chain features. For the ClpX reconstruction, we extracted ClpX sub-particles from both ends of ClpP based on the previously determined ClpP alignment using a python script and IMOD (Mastronarde and Held, 2017). We prepared an original and a ClpP-signal-subtracted ClpX particle stack. To remove misaligned particles, the ClpX stacks were 2D classified without alignment. Particles from 2D classes that showed clear secondary structure were used in subsequent 3D-classification and reconstruction steps.

Clean ClpX sub-particles were 3D classified (K=6) and refined, resulting in four distinct ClpX classes with resolutions ranging from 3.9 to 4.2 Å. To recover the ClpX^ΔN^-ClpP interface, we re-extracted ClpXP sub-particles using a larger box that included the cis ClpP ring. Alignment and classification of each ClpX^ΔN^ sub-particle was transferred to corresponding ClpX^ΔN^/ClpP sub-particle and four classes were refined with local alignment optimization, resulting in four ClpX^ΔN^/ClpP maps (resolutions 4.0 to 4.3 Å).

To test the robustness of this workflow, we performed 3D classification of ClpX sub-particles multiple times using K=4 or K=8. The quality of maps suffered slightly, but the overall structures of four predominant classes of ClpX hexamers remained unchanged.

### Model building and refinement

A ClpP tetradecamer (PDB 3MT6; Li et al., 2010) was docked into the D_7_ map using Chimera’s “fit to map” function (Pettersen et al., 2004). For ClpX^ΔN^, six copies of the large and small AAA+ domains (PDB 3HWS; Glynn et al., 2009) were docked into the map sequentially and refined for three iterations in Chimera. Real-space refinement of docked ClpP and ClpX was performed using PHENIX (Adams et al., 2010), and model building was performed using COOT (Emsley and Cowtan, 2004). ChimeraX (Goddard et al., 2014), Chimera (Pettersen et al., 2004), and PyMOL were used to create figures and movies.

### Biochemical assays

Assays were conducted at 37 °C in PD buffer (25 mM HEPES-KOH, pH 7.5, 5 mM MgCl_2_, 200 mM KCl, 10% glycerol). Experiments were performed in triplicate and reported values are averages ± SD. Degradation of ^CP7^GFP-ssrA (15 µM) by ClpX^ΔN^ (0.3 µM hexamer) and ClpP_14_ (0.9 µM) was assayed in the presence of 5 mM ATP and was monitored by loss of fluorescent signal (excitation 467 nm; emission 511 nm). For RKH loop mutants with reduced substrate affinity, degradation was measured at high concentrations of fluorescent substrate, with excitation at an off-peak wavelength (excitation 420 nm; emission 511 nm). Degradation of fluorescein-labeled Arc-st11-ssrA by ClpX^ΔN^ (0.3 µM hexamer) and ClpP_14_ (0.9 µM) was assayed as described (Bell et al., 2018). *K*_M_ and V_max_ were determined by fitting the average values of replicates to a hyperbolic equation. ATP hydrolysis rates were measured using a coupled-NADH oxidation assay as described (Martin et al., 2005), using ClpX^ΔN^ (0.3 µM hexamer), with or without ClpP_14_ (0.9 µM), and 5 mM ATP. Activation of decapeptide cleavage by ClpP or variants was performed as described (Lee et al., 2010A), using ClpP_14_ (50 nM), RseA decapeptide (15 µM), ATP (5 mM) and a regeneration system, and either ClpX^ΔN^ (0.5 µM hexamer) or ADEB-2B (100 µM), which was a generous gift from J. Sello (Brown). Pull-down experiments were performed in 25 mM HEPES-KOH (pH 7.5), 5 mM MgCl_2_, 150 mM KCl, 20 mM imidazole, 10% glycerol, 500 µM dithiothreitol and 2 mM ADP or ATPγS. 40 µL of a mixture of ClpX^ΔN^ C169S (1 µM hexamer concentration) and ClpP (1 µM 14-mer concentration) were mixed, incubated for 10 min at 30 °C, and then added to 20 µL of Ni^2+^-NTA resin (Thermo Scientific) equilibrated in the same buffer. Binding reactions were incubated for 15 min at room temperature with rotation, centrifuged for 1 min at 9400 x g, and the supernatant was discarded. Reactions were washed, centrifuged, and the supernatant discarded three times. Bound protein was eluted with 40 µL of buffer supplemented with 300 mM imidazole for 15 min with rotation. Reactions were then centrifuged again and the eluant collected. Input and elution samples for each reaction were resolved by SDS-PAGE on a 10% Bis-Tris/MES gel run at 150V and visualized by staining with Coomassie Brilliant Blue R250. Results were validated in three independent replicates.

## ACKNOWLEDGMENTS

Supported by NIH grant GM-101988 (R.T.S) and the Howard Hughes Medical Institute (T.A. Baker; S.C. Harrison). We thank S. Glynn, M. Lang, H. Manning, A. Olivares, and K. Schmitz for helpful discussions, J. Sello for providing ADEP-2B, and C. Xu and K. Song at the Electron Microscopy Facility at the University of Massachusetts Medical School for advice and data collection. The authors declare no competing financial interests.

## AUTHOR CONTRIBUTIONS

X.F., S.J., and R.T.S. performed cryo-EM experiments, calculations, and/or model building and refinement. T.A. Bell and B.M.S. performed biochemical experiments. S.P.H., T.A. Baker, and R.T.S. oversaw research. All authors contributed to writing and/or revising the manuscript.

## Supplement

**Figure S1.**
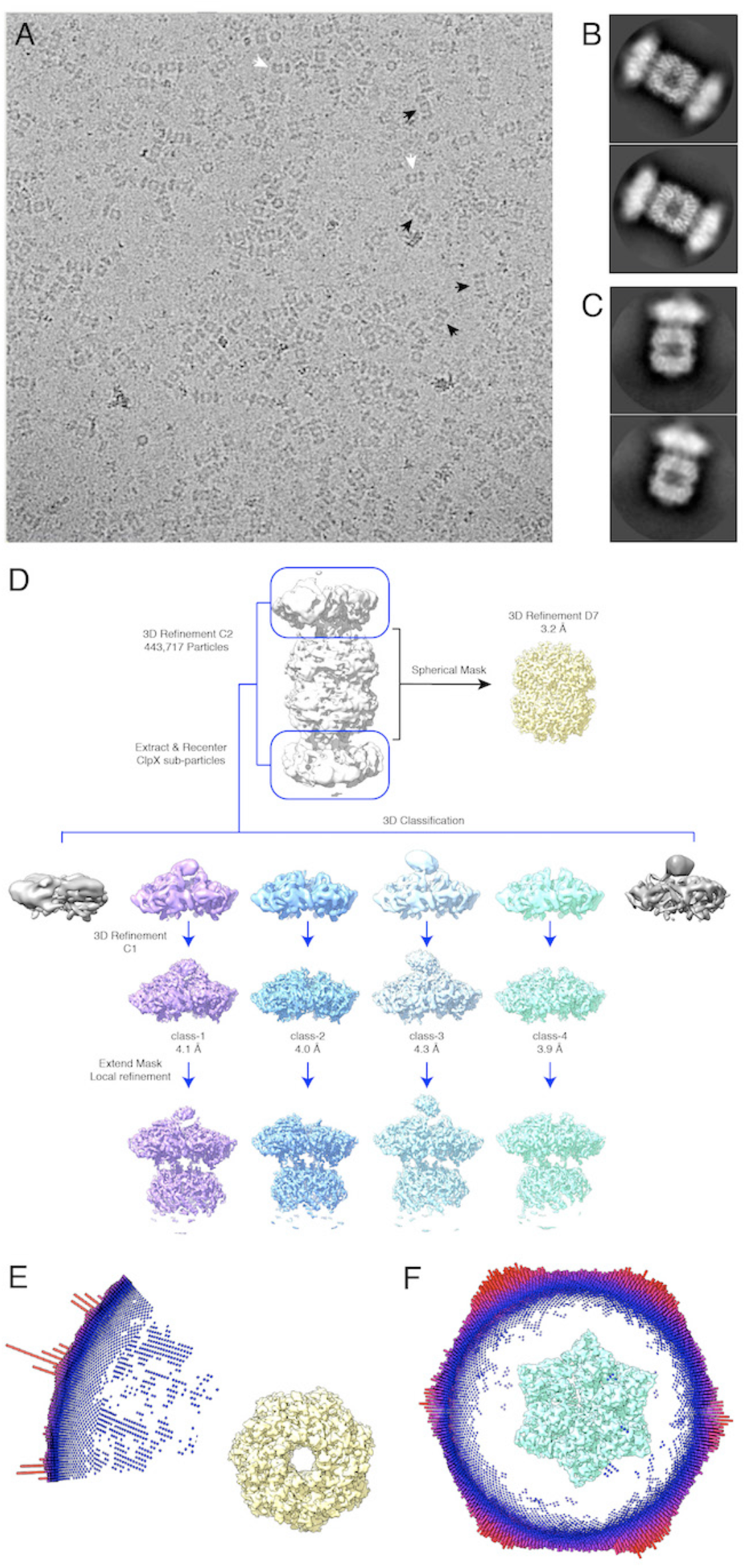
Cryo-EM data and strategy. (A) Cryo-EM micrograph at 36000X magnification of doubly capped particles (black arrows) and singly capped particles (white arrows). (B) 2D-class averages of doubly capped complexes. (C) 2D-class averages of singly capped complexes. (D) Relion data processing scheme used to obtain 3D reconstructions. (E) Euler-angle distribution of particles used to reconstruct the D_7_ symmetric ClpP map. (F) Euler-angle distribution of particles used to reconstruct the non-symmetric class-4 ClpX^ΔN^ map.

**Figure S2.**
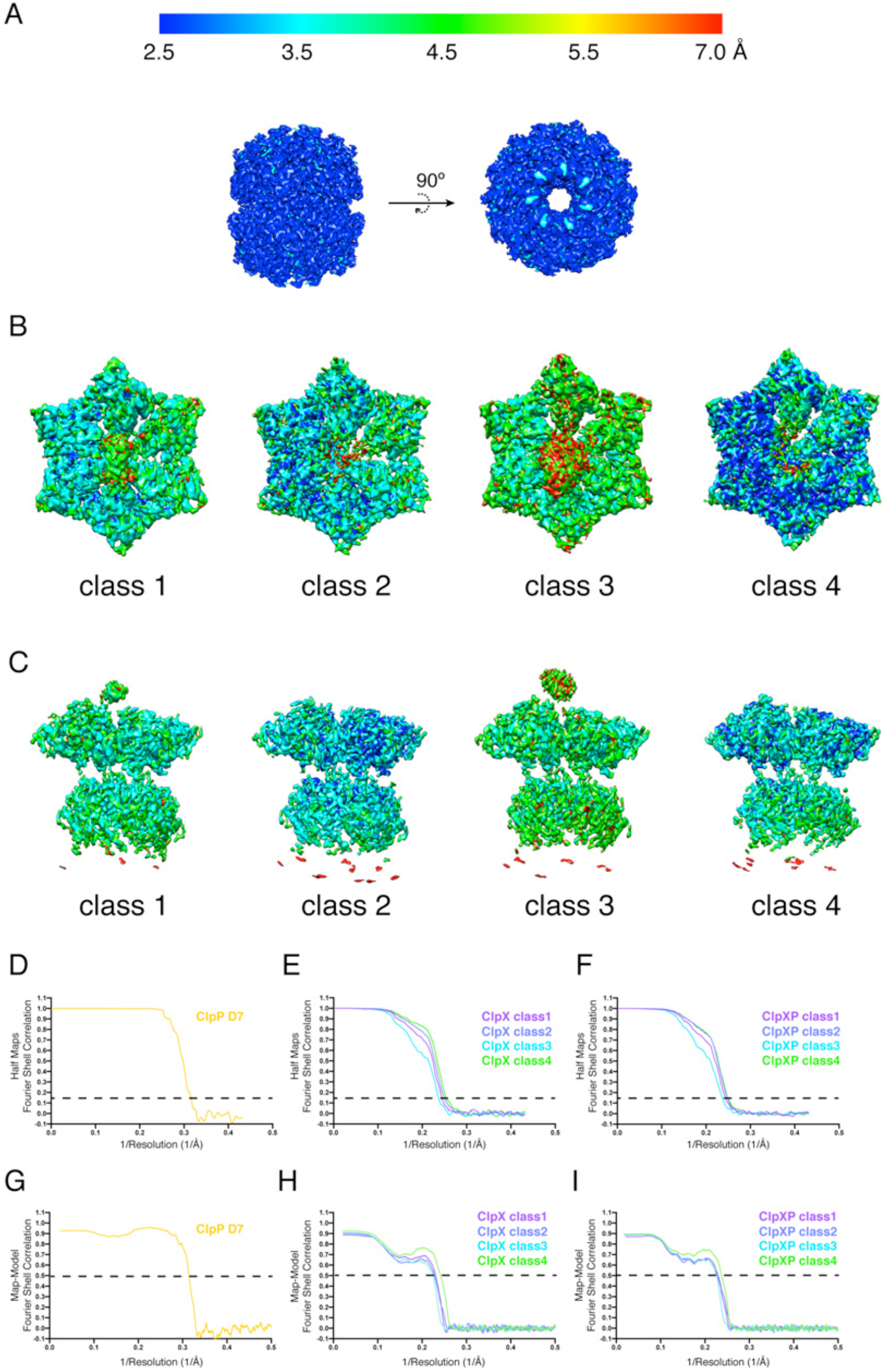
Local map resolution and FSC plots. (A) Side views and top views of ClpP (D_7_). (B) Top views of the four ClpX^ΔN^ classes. (C) Side views of the four ClpXP classes. ResMap^61^ was used to estimate local resolutions. (D-F) FSC plots of all masked final 3D reconstructions. (G-I) FSC plots calculated using masked 3D reconstructions and models.

**Figure S3.**
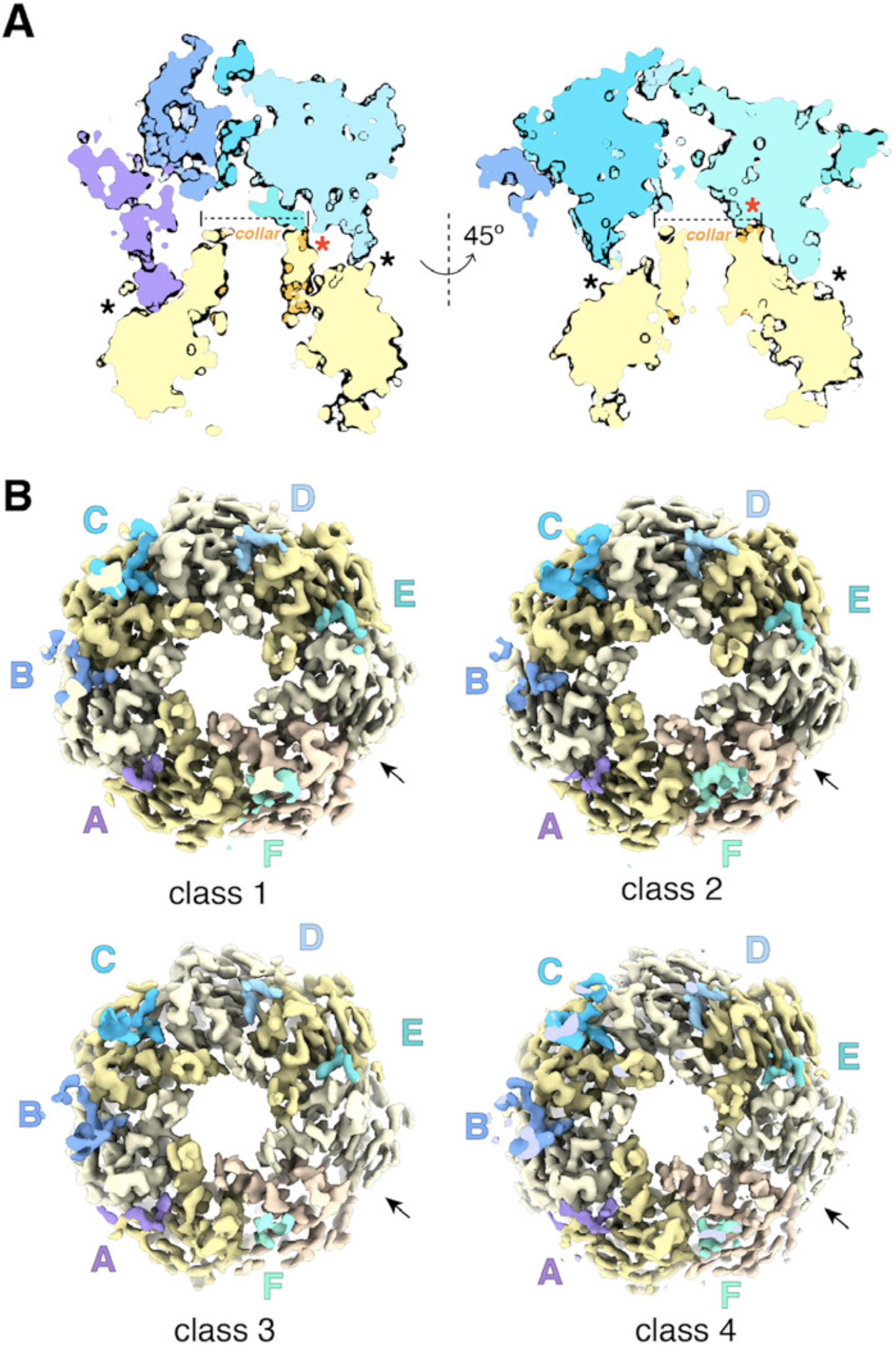
Structural features of ClpX^ΔN^/ClpP interface. (A) Cutaway showing axial collar of ClpP fitting into a bowl-shaped cavity at the bottom of the ClpX^ΔN^ hexamer. Black asterisks show contacts between the ClpX IGF loops and ClpP clefts. Red asterisks show contacts between ClpX and the ClpP collar. (B) Top views of ClpXP classes with substrate and most ClpX^ΔN^ except IGF loops (blue or purple) removed. Arrows show the position of the empty ClpP cleft without an IGF loop.

**Figure S4.**
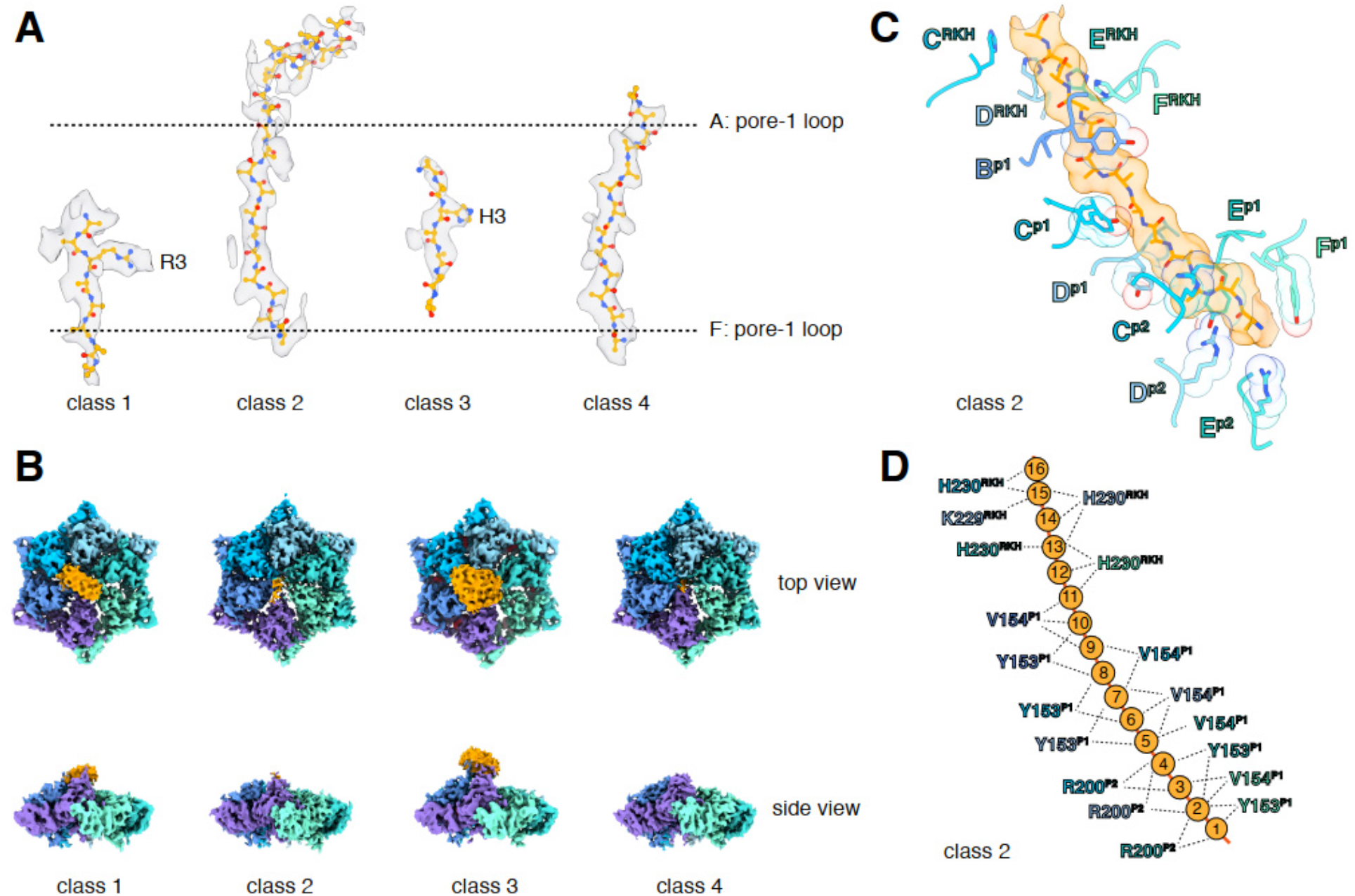
Substrate interactions with ClpX. (A) Substrate density within the ClpX channel is rendered as a transparent surface (grey) with atoms shown in ball and stick representation. Dashed lines shown approximate positions of the pore-1 loop from the ClpX subunit A (top) and subunit F (bottom). (B) Substrate density above the ClpX hexamer. Substrate density is colored in orange and ClpX subunits are colored in different shades from purple to aquamarine. (C) Interactions between substrate (orange, stick and surface representation) and ClpX pore loops (blue and purple, stick and transparent space-filling representation) in the class-2 structure. Capital letters indicate ClpX subunits; superscripts indicate RKH loops, pore-1 (p1) loops, or pore-2 (p2) loops. (D) Scheme of interactions shown in panel C. Dashed lines represent distances of 6.5 Å or less between the Cβ atoms of substrate alanines and the Cβ atoms of Y153/V154 (p1), or the Cε atom of R200 (p2), or the Cγ atom of H230 (RKH), or the Nζ atom of R228 (RKH).

**Figure S5.**
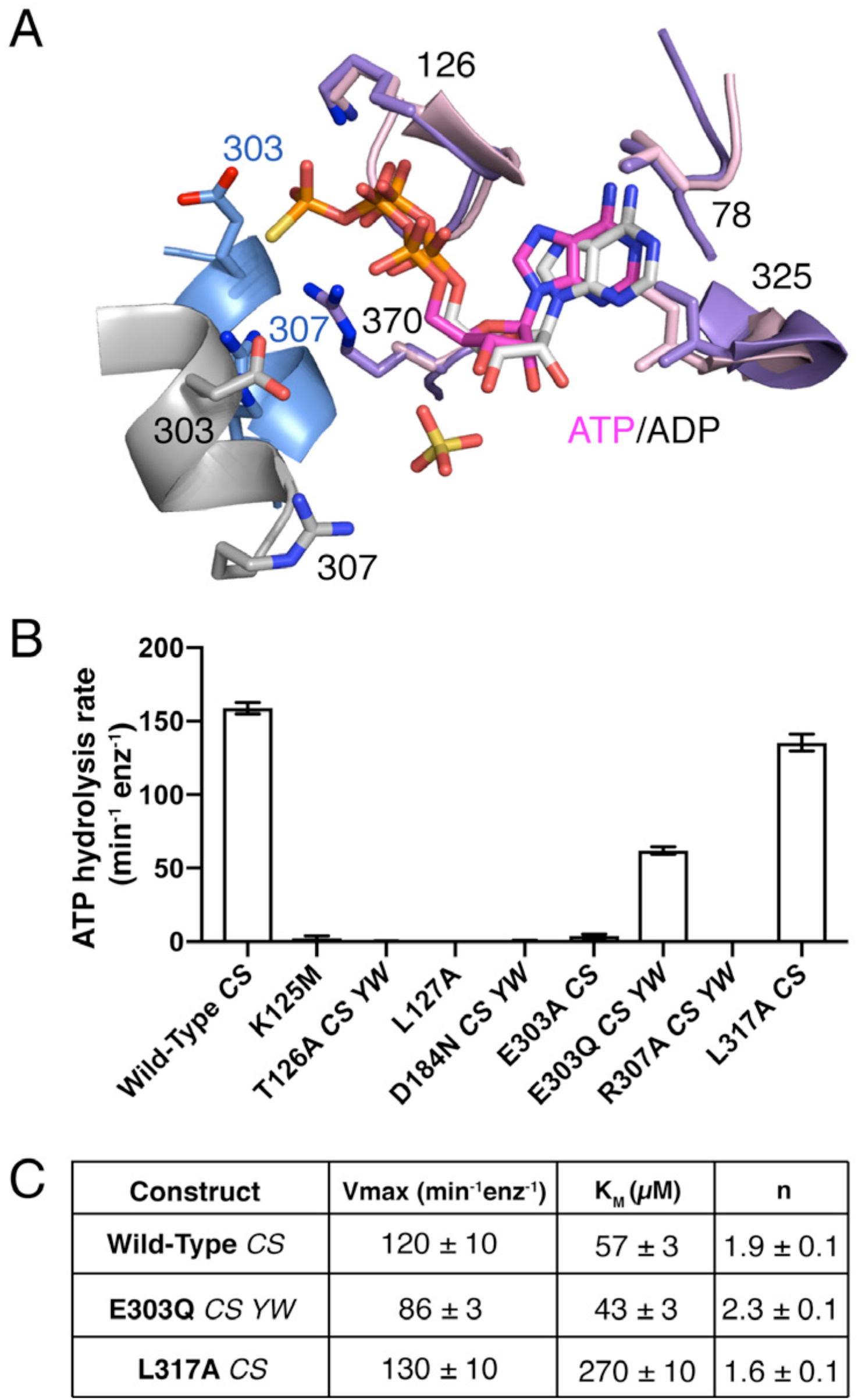
Nucleotide-binding pocket and mutations. (A) Alignment of residues contacting bound nucleotide in the class-4 cryo-EM structure (A/B subunit interface; purple/blue) and crystal structure of ClpX^ΔN^ (pink/grey; PDB 3HWS; Glynn et al., 2009). (B) ATP-hydrolysis activity for ClpX^ΔN^ variants with mutations in nucleotide-binding pocket. Values are averages of three independent replicates ± SD. (C) Fitted Michealis-Menten-Hill parameters for ATP hydrolysis for variants with significant activity. Values are averages of three independent replicates ± SD. In panels B and C, *CS* (C169S) and *YW* (Y77W) indicate additional mutations in variants that do not, by themselves, alter ATP hydrolysis activity.

**Figure S6.**
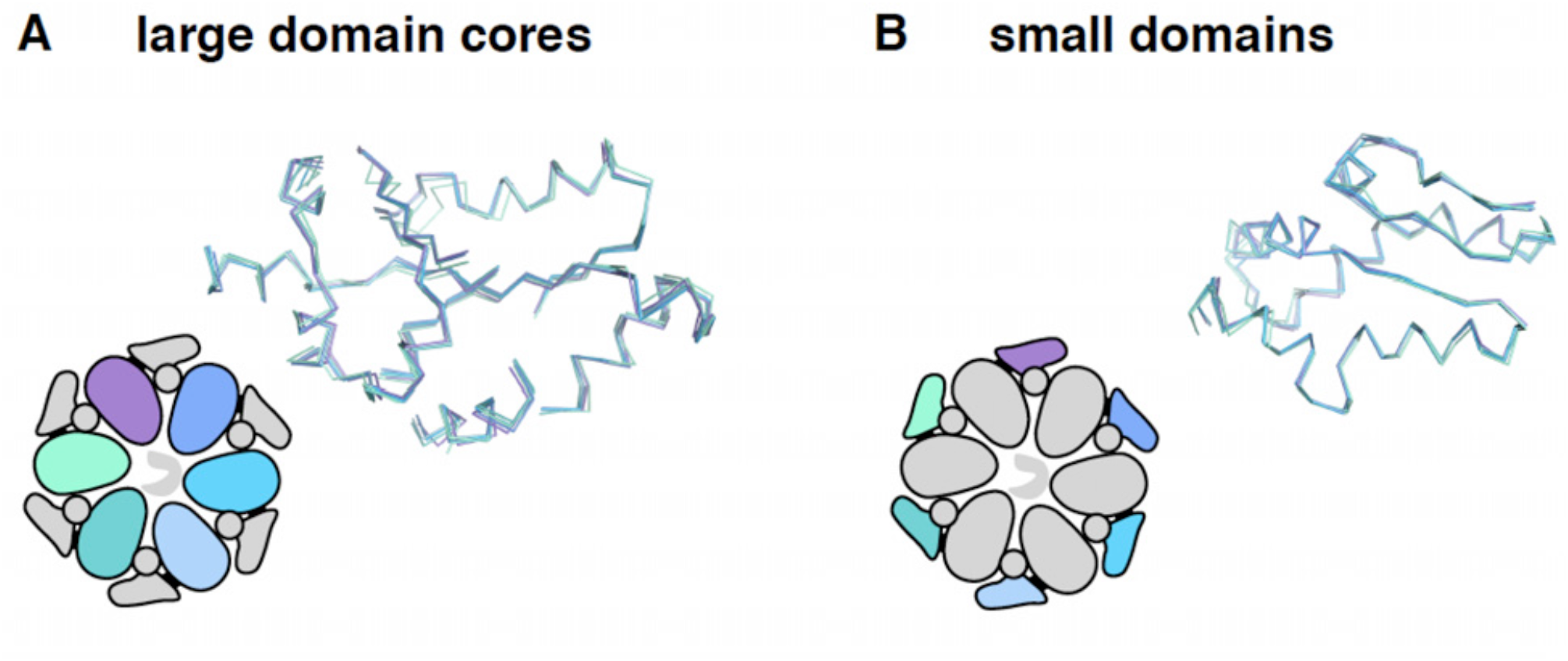
Invariant and flexible portions of ClpX^ΔN^. (A) Superimposition of the cores of the large AAA+ domains from each subunit of the class-4 ClpX^ΔN^ structure. (B) Superimposition of the small AAA+ domains from the class-4 ClpX^ΔN^ hexamer.

**Figure S7.**
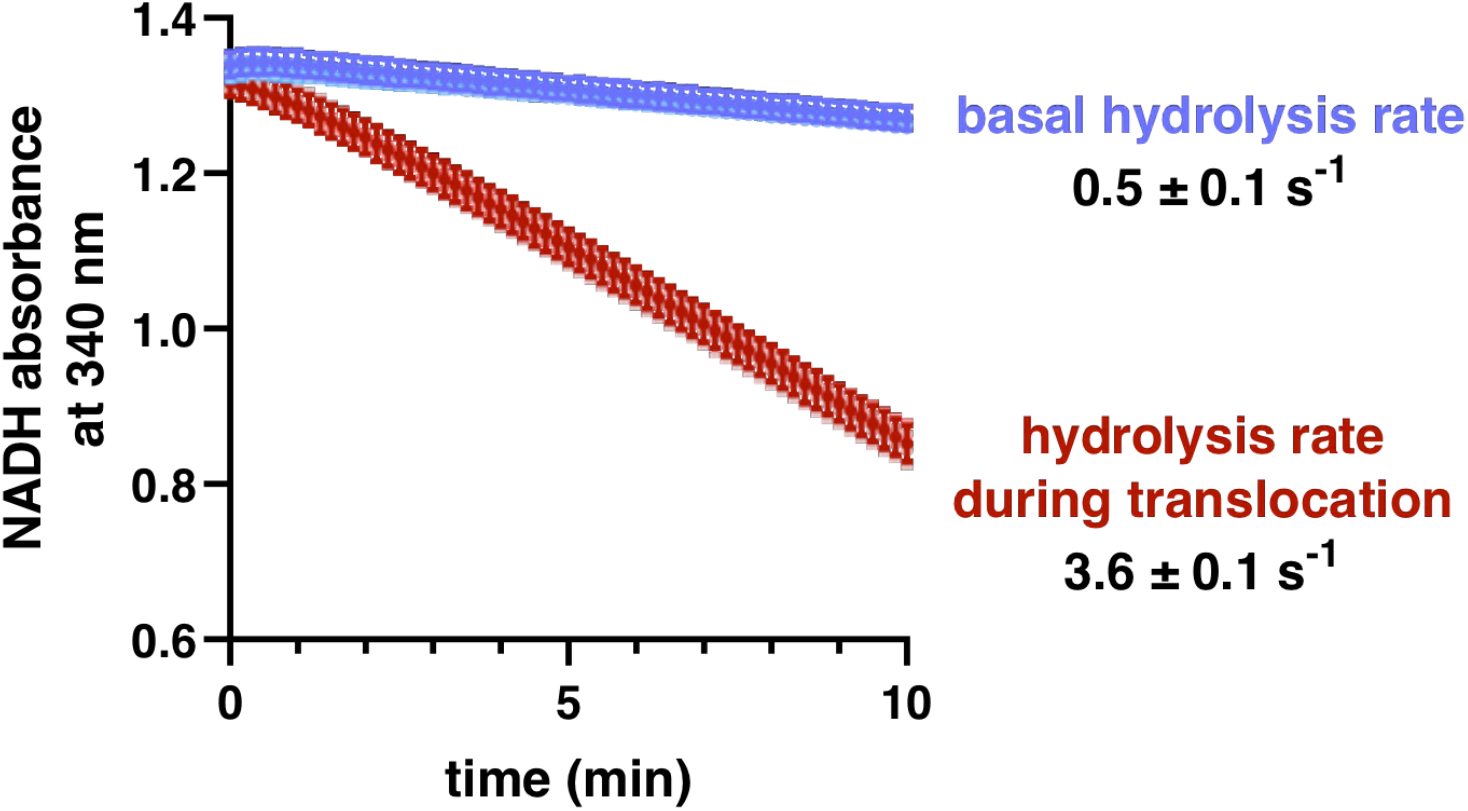
ATP-hydrolysis rates. ATP-hydrolysis rates by ClpX^ΔN^/ClpP were measured using a coupled assay (Martin et al., 2005) that results in loss of NADH absorbance at 23 °C in PD buffer, conditions that approximate the conditions of optical-trapping experiments (Cordova et al., 2014). Shown are experiments in the absence (basal) or presence (translocation) of a near-saturating concentration (20 µM) of an unfolded titin^I27^-ssrA substrate. The figure shows an overlay of three independent replicates. The rates shown are averages of the three independent replicates ± SD in units corresponding ATP molecules hydrolyzed per second per enzyme.

**Table S1.** Cryo-EM data collection, processing, model building and validation statistics.

| Name | ClpP | ClpX-class 1 | ClpX-class 2 | ClpX-class 3 | ClpX-class 4 | ClpXP-class 1 | ClpXP-class 2 | ClpXP-class 3 | ClpXP-class 4 |
| --- | --- | --- | --- | --- | --- | --- | --- | --- | --- |
| **PDB ID** | 6PPE | 6PP8 | 6PP7 | 6PP6 | 6PP5 | 6POS | 6POD | 6PO3 | 6PO1 |
| **EMDB ID** | 20434 | 20422 | 20421 | 20420 | 20419 | 20418 | 20412 | 20408 | 20406 |
| **Data collection and processing** |  |  |  |  |  |  |  |  |  |
| Microscope | Talos Arctica |  |  |  |  |  |  |  |  |
| Camera | K2 summit |  |  |  |  |  |  |  |  |
| Magnification | 36000X |  |  |  |  |  |  |  |  |
| Voltage (kV) | 200 |  |  |  |  |  |  |  |  |
| Total electron dose (e-/Å<sup>2</sup>) | 58 |  |  |  |  |  |  |  |  |
| Defocus range (µm) | -1.2 to -2.5 |  |  |  |  |  |  |  |  |
| Pixel size (Å) | 0.58 |  |  |  |  |  |  |  |  |
| Micrographs collected | 3657 |  |  |  |  |  |  |  |  |
| Final particles | 443717 | 151652 | 241899 | 115751 | 227257 | 151652 | 241899 | 115751 | 227257 |
| Symmetry | D<sub>7</sub> | C<sub>1</sub> | C<sub>1</sub> | C<sub>1</sub> | C<sub>1</sub> | C<sub>1</sub> | C<sub>1</sub> | C<sub>1</sub> | C<sub>1</sub> |
| Resolution (FSC 0.143) | 3.19 | 4.12 | 4.05 | 4.28 | 3.98 | 4.12 | 4.05 | 4.28 | 4.05 |
| **Model composition** |  |  |  |  |  |  |  |  |  |
| Non-hydrogen atoms | 21840 | 15498 | 15593 | 15599 | 15530 | 26656 | 26626 | 26597 | 26336 |
| Protein residues | 2814 | 1994 | 2012 | 2006 | 2000 | 3432 | 3431 | 3425 | 3396 |
| Ligands | 0 | 6 | 6 | 6 | 6 | 6 | 6 | 6 | 6 |
| **Refinement** |  |  |  |  |  |  |  |  |  |
| Map-model CC | 0.90 | 0.76 | 0.75 | 0.76 | 0.8 | 0.76 | 0.73 | 0.72 | 0.77 |
| Map sharpening B factors (Å<sup>2</sup>) | -180 | -186 | -235 | -216 | -170 | -160 | -176 | -184 | -184 |
| R.m.s deviations - bond length (Å) | 0.004 | 0.002 | 0.003 | 0.003 | 0.003 | 0.002 | 0.002 | 0.002 | 0.002 |
| R.m.s deviations - bond angles (degrees) | 0.648 | 0.637 | 0.667 | 0.664 | 1.164 | 0.650 | 0.666 | 0.658 | 0.660 |
| **Validation** |  |  |  |  |  |  |  |  |  |
| MolProbity score | 0.60 | 0.94 | 0.94 | 0.94 | 0.91 | 0.91 | 0.83 | 0.91 | 0.90 |
| Clashscore | 0.27 | 1.83 | 1.78 | 1.78 | 1.63 | 1.60 | 1.16 | 1.63 | 1.53 |
| C-beta deviations | 0 | 0 | 0 | 0 | 0 | 0 | 0 | 0 | 0 |
| Rotamers outlier (%) | 0 | 0 | 0.18 | 0 | 0 | 0 | 0.04 | 0 | 0 |
| Ramachandran favored (%) | 98.51 | 99.95 | 99.85 | 99.90 | 99.85 | 99.91 | 99.76 | 99.73 | 99.97 |
| Ramachandran allowed (%) | 1.49 | 0.05 | 0.15 | 0.10 | 0.15 | 0.09 | 0.24 | 0.27 | 0.03 |
| Ramachandran disallowed (%) | 0 | 0 | 0 | 0 | 0 | 0 | 0 | 0 | 0 |

**Table S2.** Model to map correlation coefficient (CC) of amino acids and nucleotides. Calculated using phenix_realspace.correlation (Adams et al., 2010). Nucleotide sites with suspected partial occupancy are marked in bold.

|  |  | **A** | **B** | **C** | **D** | **E** | **F** |
| --- | --- | --- | --- | --- | --- | --- | --- |
| **class 1** | protein | 0.78 | 0.79 | 0.81 | 0.81 | 0.81 | 0.80 |
|  | nucleotide | 0.77 | 0.80 | 0.82 | 0.81 | 0.84 | 0.74 |
| **class 2** | protein | 0.81 | 0.83 | 0.82 | 0.82 | 0.80 | 0.80 |
|  | nucleotide | **0.59** | 0.87 | 0.80 | 0.80 | 0.81 | **0.67** |
| **class 3** | protein | 0.82 | 0.83 | 0.80 | 0.80 | 0.80 | 0.78 |
|  | nucleotide | 0.79 | 0.84 | 0.83 | 0.80 | 0.77 | 0.76 |
| **class 4** | protein | 0.81 | 0.76 | 0.74 | 0.73 | 0.76 | 0.80 |
|  | nucleotide | 0.83 | 0.77 | 0.78 | 0.74 | 0.78 | **0.65** |

**Table S3.**
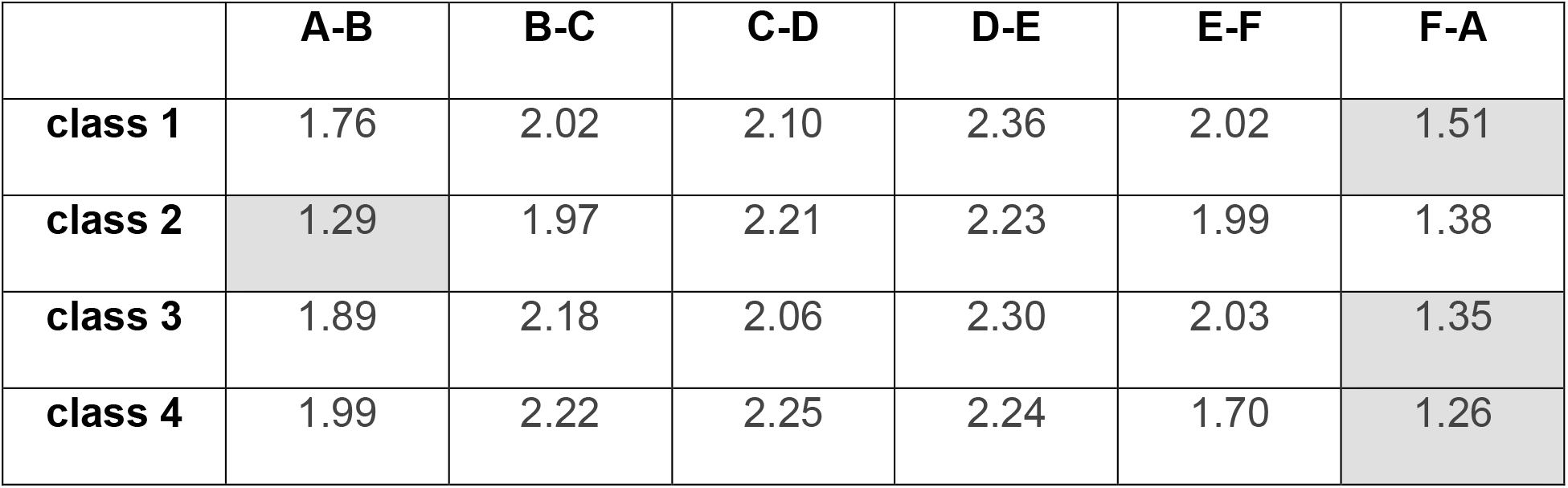
Buried surface area (in units of 1000 Å^2^) between ClpX^ΔN^ subunits. The interface in each hexamer with the smallest amount of buried surface is highlighted in gray and corresponds to the ADP-bound subunit.

|  | **A-B** | **B-C** | **C-D** | **D-E** | **E-F** | **F-A** |
| --- | --- | --- | --- | --- | --- | --- |
| **class 1** | 1.76 | 2.02 | 2.10 | 2.36 | 2.02 | 1.51 |
| **class 2** | 1.29 | 1.97 | 2.21 | 2.23 | 1.99 | 1.38 |
| **class 3** | 1.89 | 2.18 | 2.06 | 2.30 | 2.03 | 1.35 |
| **class 4** | 1.99 | 2.22 | 2.25 | 2.24 | 1.70 | 1.26 |

**Table S4.** Distances between nucleotides and key residues. ADP-occupied subunits are highlighted in gray. Measurements in Å are between the γ thiophosphate of ATPγS or β phosphate of ADP and the Cε of arginine or Cγ of glutamine.

| **class 1** | **R307**<br>**(Arg finger)** | **R370**<br>**(sensor II)** | **E185**<br>**(Walker B)** |
| --- | --- | --- | --- |
| A | 6.2 | 4.2 | 7.1 |
| B | 4.5 | 4.6 | 7.0 |
| C | 4.7 | 4.7 | 6.2 |
| D | 4.8 | 4.4 | 6.2 |
| E | 5.7 | 4.6 | 6.1 |
| F | 15.1 | 4.7 | 8.8 |
| **class 2** | **R307** | **R370** | **E185** |
| A | 8.0 | 4.0 | 7.9 |
| B | 4.9 | 4.0 | 6.8 |
| C | 5.2 | 4.2 | 6.0 |
| D | 4.4 | 4.3 | 6.6 |
| E | 5.9 | 4.5 | 6.6 |
| F | 16.2 | 4.6 | 6.3 |
| **class 3** | **R307** | **R370** | **E185** |
| A | 5.6 | 4.2 | 6.7 |
| B | 4.6 | 4.3 | 6.7 |
| C | 5.4 | 4.2 | 6.5 |
| D | 4.7 | 4.6 | 5.9 |
| E | 5.6 | 6.3 | 6.4 |
| F | 15.9 | 5.8 | 5.8 |
| **class 4** | **R307** | **R370** | **E185** |
| A | 6.2 | 4.0 | 7.0 |
| B | 4.6 | 4.3 | 6.6 |
| C | 4.7 | 4.6 | 6.9 |
| D | 4.6 | 4.3 | 6.4 |
| E | 8.8 | 5.1 | 5.9 |
| F | 13.8 | 4.6 | 7.7 |

## Movie Legends

Movie S1. Top and side views of a ClpX^ΔN^ hexamer and adjacent ClpP ring for classes 1, 2, 3, and 4. ClpX^ΔN^ subunits (cartoon representation) are colored blue, green, or purple; carbons in substrate (space-filling representation) are colored orange; carbons in ATP or ADP (space-filling representation) are colored magenta; and ClpP subunits (cartoon representation) are colored yellow with the axial β-hairpin colored gold. Note that substrate above the axial channel of ClpX^ΔN^ is not shown because the corresponding density was not modeled.

Movie S2. Interaction of substrate with ClpX^ΔN^. Substrate in the axial channel of the class-4 ClpX structure adopts an extended conformation and is contacted by the pore-1, pore-2, and RKH loops of ClpX, which are arranged into a spiral and pack against each other.

Movie S3. Transition between class-2 and class-4 structures of ClpX^ΔN^/ClpP. A conformational morph between the class-2 and class-4 structures of ClpXP is shown. To generate this morph, ClpX subunits in the class-2 structure were renamed from ABCDEF to FABCDE and aligned to the class-4 ClpXP structure in two steps to maximize overlap between the ClpP portions of the two structures. The ADP-bound ClpX subunit (colored aquamarine) moves most dramatically, from the top to the bottom of the ClpX spiral. The last visible residue of the pore-1 loop of the ADP-bound subunit (T149) is colored red and moves ∼2.7 nm.

